# Coordinated multi-level adaptations across neocortical areas during task learning

**DOI:** 10.1101/2024.09.26.615162

**Authors:** Shuting Han, Fritjof Helmchen

## Abstract

The coordinated changes of neural activity during learning, from single neurons to populations of neurons and their interactions across brain areas, remain poorly understood. To reveal specific learning-related changes, we applied multi-area two-photon calcium imaging in mouse neocortex during training of a sensory discrimination task. We uncovered coordinated adaptations in primary somatosensory area S1 and the anterior (A) and rostrolateral (RL) areas of posterior parietal cortex (PPC). At the single-neuron level, task-learning was marked by increased number and stabilized responses of task neurons. At the population level, responses exhibited increased dimensionality and reduced trial-to-trial variability, paralleled by enhanced encoding of task information. The PPC areas, especially area A, became gradually engaged, opening additional within-area subspaces and inter-area subspaces with S1. Task encoding subspaces gradually aligned with these interaction subspaces. Behavioral errors correlated with reduced neuronal responses, decreased encoding accuracy, and misaligned subspaces. Thus, multi-level adaptations within and across cortical areas contribute to learning-related refinement of sensory processing and decision-making.

## Introduction

Learning a new task requires accurate sensory processing and transforming relevant sensory information into appropriate actions. This process often involves not only reconfigurations of local neuronal populations, but also coordinated changes across brain areas. Many studies have characterized learning-related changes at single-neuron and population coding levels within local populations^1–7^. These changes include increased stimulus selectivity^1,6^, recruitment of new neurons^3,6^, stabilized responses of individual neuron^2,3,7^, and increased task information encoded by the populations^1–6^. Recently, an increasing number of studies focused on understanding how population activity dynamics, which often reflects task-related information, changes with learning^8–12^. For example, learning a new task is accompanied by stabilized population dynamics, and expanded activity subspaces of intrinsic dynamic to encode task-relevant variables^10,13–15^. The local population dynamics can also constrain the learning process: learning something new within the intrinsic subspace of neural dynamics is easier than outside of the subspace^16^, and the internal state of the animal, e.g., the level of task engagement, can shape learning through changing these subspaces^17^.

Understanding such population-level changes is important for unveiling the neural computations underlying the optimization of sensory processing during learning, supported by changes on the single-neuron level.

Beyond the changes in local populations, task-learning is often accompanied by coordinated changes across brain areas. Learning to perform a specific task typically involves a distinct set of brain areas, with specific higher areas becoming increasingly engaged to selectively process task-relevant information sent by primary sensory areas^6,18,19^. However, we still poorly understand how information processing is transformed in a coordinated way across multiple areas during learning, from primary sensory areas that receive sensory inputs, to higher areas that further process sensory information, make decisions, and generate actions. Despite the rising interest in understanding cross-areal processing from the perspective of behavior-relevant subspaces as well as interaction subspaces^20–22^, it remains unclear whether within-area and inter-area interaction subspaces are reconfigured during learning to facilitate task information processing, or whether these intrinsic subspaces rather impose constraints on learning. One major challenge to answer these questions is to record from enough neurons simultaneously across multiple areas. Here, we addressed this challenge by employing a custom multi-area two-photon microscope^23,24^ to measure the learning-related changes in population dynamics, within and between areas along the cortical hierarchy, and to analyze the changes in their interaction subspaces.

A key area in processing and routing sensory information is the posterior parietal cortex (PPC). The PPC has dense connections with primary sensory regions, including the visual (V1), somatosensory (S1), and auditory (A1) cortices, as well as with frontal areas and the thalamus^25^. PPC supports a wide range of functions, such as multisensory integration, evidence accumulation, decision-making, and working memory^26–29^. Although PPC overall receives inputs from all primary sensory regions, recent works have supported the existence of functional subregions of PPC in the mouse brain: the rostrolateral region (PPC-RL) receives more inputs from V1 and S1, and is involved in visual and tactile processing, and the anterior region (PPC-A) receives more inputs from V1 and A1, and is involved in visual and auditory processing^23,26^. In particular, PPC emerges as a key area for routing sensory information during learning a texture discrimination task^26,28,30^, and is dynamically reorganized over the learning process^8^. These features make PPC an attractive example area to study the principles of learning mechanisms within and across areas.

Here we characterized learning-related changes in the local populations of whisker-related S1 barrel cortex, and PPC-RL and PPC-A. Specifically, we trained mice to perform a two-alternative auditory-cued texture discrimination task, while simultaneously imaging population activity in S1 and PPC throughout learning. We systematically examined four types of population subspaces: the variance subspace, the encoding subspace, the within-area interaction subspace, and the inter-area interaction subspace. We found that task-learning was accompanied by systematic and coordinated changes within and across these areas, both on the single-neuron level and on the population level. The PPC areas, particularly PPC-A, were gradually engaged during the learning process, expanding their within-area and inter-area interaction subspaces with S1. Task representation was improved through an increased alignment of encoding subspaces with these interaction subspaces. These learning-related increases were absent during incorrect trials. These results suggest that refinement of sensory and choice processing during learning is achieved through coordinated adaptations on multiple levels across neocortical areas, from enhanced responses of task neurons, to improved alignment of task-relevant information to the intrinsic dynamic subspaces within and across areas.

## Results

### Behavioral task

We trained 16 mice (age 2-4 months, both sexes) expressing GCaMP6f in L2/3 to perform an auditory-cued two-alternative texture discrimination task, while monitoring neuronal population activity in S1 and PPC^23^ (Figure 1a; Methods). The task consisted of four sequential phases (Figure 1b; Methods): each trial started with an auditory tone (‘tone window’), followed by presentation of sandpaper texture to the whisker pad (‘texture window’). A low tone was always paired with a rough sandpaper, and a high tone with a smooth sandpaper. Then, the mouse chose between two lick ports (‘choice window’) to obtain a sugar water reward (‘reward window’). The reward window was triggered as soon as mice had made their choice. Each day, mice performed between 100 and 500 trials during training. To account for the different trial numbers each day, we split each training day into sessions of 80-120 trials. To study learning-related changes, we defined three learning phases according to the behavioral performance: naive (performance<55%), learning (performance 55%-75%), and expert (performance ≥75%) phase (Fig. 1c). During the expert phase, a subset of the sessions included a small fraction (10-30%) of tone-texture mismatch trials, in which the tone was paired with the non-matching texture, and reward was given according to texture. These data revealed interesting results about predictive processing and have been previously published^23^. Due to the low percentage of mismatch trials and because we did not observe re-learning due to these mismatch trials, we included this expert dataset for analysis, but excluded all the mismatch trials, as well as the sessions with low behavioral performance (<70%).

**Figure 1.**
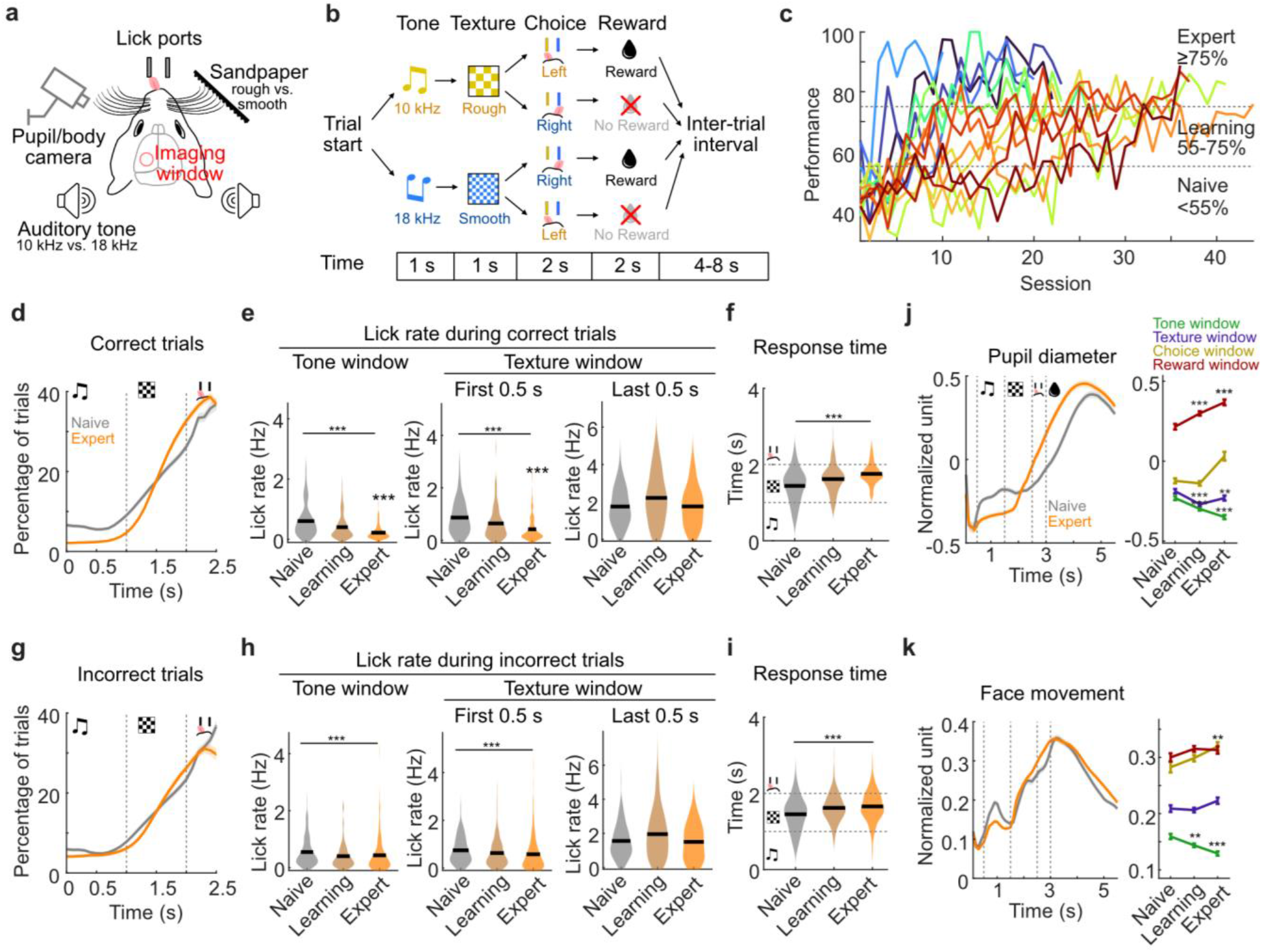
Learning behavior of mice in a two-alternative tone-texture discrimination task. **(a)** Schematic of Experimental setup. **(b)** Task design. Each trial consisted of a tone followed by a paired texture presented to the whisker pad of the mice. Then, mice chose between two lick ports to receive a sugar water drop as reward. **(c)** Learning curves of all mice (n = 16), subdivided into sessions of 80-120 trials. Naive is defined as performance <55%, learning as 55-75%, and expert as ≥75%. Different colors indicate mice. **(d)** Lick pattern in correct trials, represented as percentage of trials with a lick at each time point, in naïve (grey line) and expert (orange line) mice. **(e)** Quantification of lick rates during tone window (left), early texture window (middle), and late texture window (right), in correct trials (black line represents mean). **(f)** Response time in correct trials. **(g)** Lick pattern in incorrect trials, represented as percentage of trials with lick at each time point, in naive and expert mice. **(h)** Quantification of lick rates during tone window (left), early texture window (middle), and late texture window (right), in incorrect trials. **(i)** Response time in incorrect trials. **(j)** Pupil diameter (z-scored) over trial time (left) and quantification over learning (right). Gray and orange in the left panel indicate naive and expert condition; colors in the right panel indicate task windows. **(k)** Face movement (z-scored) over trial time (left) and quantification over learning (right). Color as in **j**. (16 mice, naive 138 sessions, learning 192 sessions, expert 161 sessions; mean±SEM)

Task-learning was accompanied by various behavioral changes. Compared to naive sessions, mice showed reduced licking in the tone and early texture windows in expert sessions (Fig. 1d-e), as well as reduced early responses (Fig. 1f). These changes were present in both correct and incorrect trials (Fig. 1d-i), suggesting a systematic change of the behavioral strategy of mice. We also monitored the pupil and movement of mice with a behavior camera (Fig. 1a). The pupil diameter during the reward window was larger in expert compared to naive sessions (Fig. 1j), and face movements were reduced during the tone window, consistent with the reduced licking behavior (Fig. 1k). These observations suggest a refinement of the task-related behavior of the mice over the learning process.

### Recruitment of task neurons in S1 and PPC during learning

To study learning-related changes across the neocortical areas, we used a custom-built two-area two-photon microscope to simultaneously record from neuronal populations in S1 barrel cortex and the two PPC subregions^23,24^. In our experiments, we simultaneously imaged somatic calcium signals from S1 and PPC-RL (total 13 mice) or from S1 and PPC-A (total 10 mice), from 2-3 depths (100-300 µm) in L2/3, covering 50-600 neurons in each area (Fig. 2a; Methods). The exact locations of S1, PPC-RL and PPC-A were identified by sensory mapping and retinotopic mapping (Methods). We extracted ΔF/F traces as well as the deconvolved spike rates of individual neurons using Suite2p^31^, and all following analysis concerning neuronal activity was performed on the z-scored spike rates. Over the course of training, we could follow roughly the same field of views (FOVs) with similar imaging quality (Fig. 2b-c). However, due to restrictions by the dense labeling in the GCaMP6f transgenic mouse lines, the large FOVs across multiple depths, and the long training process, we did not seek to match and track individual neurons across days for each mouse.

**Figure 2.**
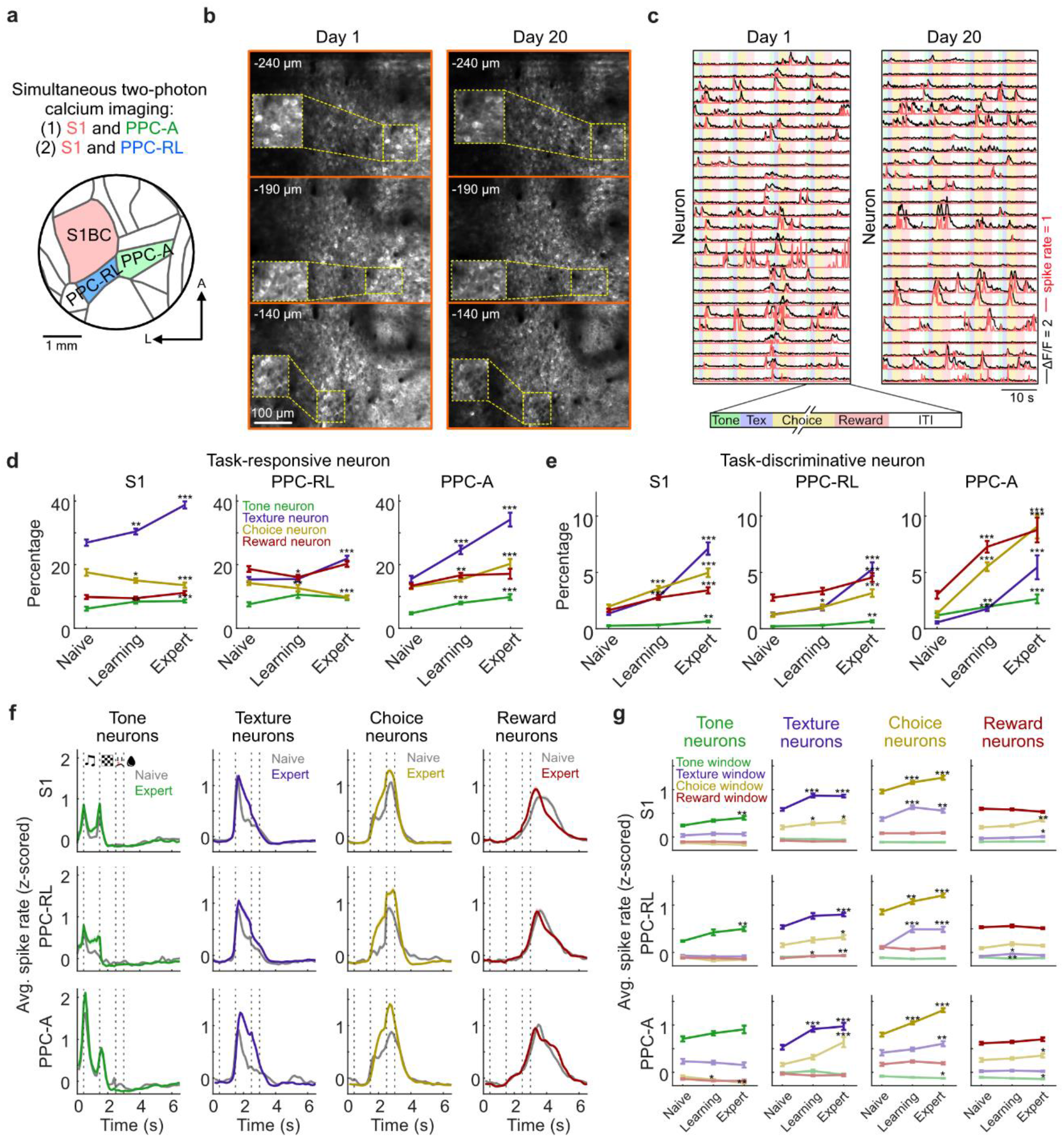
Task neurons in S1 and PPC areas. **(a)** Diagram of the cranial window and S1/PPC locations. Simultaneous two-photon calcium imaging was performed in S1 and PPC areas in GCaMP6f transgenic mice. **(b)** Example FOVs at 3 imaging depths from day 1 and day 20 of the same mouse, same area (PPC-RL). **(c)** Example neuronal activity from day 1 and day 20, from the same mouse and same area as shown in **b**. Colored stripes in the background represent task windows. Black lines represent ΔF/F traces, red lines represent deconvolved spike rates in arbitrary units. **(d)** Percentage of task-responsive neurons in S1, PPC-RL, and PPC-A. Colors represent the task window to which the neurons are responsive. **(e)** Percentage of task-discriminative neurons in S1, PPC-RL, and PPC-A. Colors represent the task window to which the neurons are responsive. **(f)** Session-average spike rates of task-discriminative neurons over trial time. Spike rates were z-scored within each session. Gray lines represent naive condition, colored lines represent expert condition. Dashed vertical lines indicate task windows. **(g)** Quantification of average spike rates of task-discriminative neurons, in different task windows. Colors represent the quantified task window. (S1: 13 mice, 130, 188, 149 sessions [naive, learning, expert]; PPC-RL: 13 mice, 70, 77, 104 sessions; PPC-A: 10 mice, 60, 111, 45 sessions; \**p*<0.05, \*\**p*<0.01, \*\*\**p*<0.001; Wilcoxon rank-sum test against naive condition; mean±SEM)

We first examined whether PPC-RL and PPC-A are required for performing the task using optogenetic inhibition. We trained 8 mice expressing channelrhodopsin-2 (ChR2) in GABAergic neurons (VGAT-ChR2-EYFP transgenic line) to perform the task. Once they had reached the expert phase, we tested their performance when PPC areas were photoinhibited through activation of GABAergic neurons (Supplemental Fig. 1a). Inhibition of either PPC-RL or PPC-A during the texture window reduced behavioral performance (Supplemental Fig. 1b), suggesting that both areas are required for optimally performing the task. Additionally, we also inhibited PPC areas in tone-only and texture-only conditions, where only one stimulus modality was presented to the mice. Inhibition of either PPC-RL or PPC-A reduced mouse performance in tone-only condition (Supplemental Fig. 1c), suggesting that both areas are involved in tone-texture association. Inhibiting PPC-RL, but not PPC-A, also impaired texture-only performance (Supplemental Fig. 1d), suggesting that PPC-RL is more involved in processing pure texture information. These results indicate that both PPC areas were required for transforming sensory information adequately into correct actions in this task.

Task-learning can be accompanied by activity changes in neurons within local populations. While some studies have reported an increased number and enhanced responses of neurons with task-related activity (‘task neurons’) during learning in the neocortex, others have found the opposite^1,6,7,18^. Therefore, we first characterized the changes in task neurons in S1 and PPC over the learning process. We identified two types of task neurons: ‘task-responsive’ neurons were defined as neurons that showed significantly higher activity in specific task windows, compared with randomly sampled activity from matching number of frames outside of the window; ‘task-discriminative’ neurons were defined as a subset of responsive neurons that showed a significant difference in activity with respect to the relevant task variable in the specific window (tone 1 vs. tone 2; texture 1 vs. texture 2; lick left vs. lick right; reward left vs. reward right). These neurons were identified for each task window and the corresponding task variables, separately. Since early licks during texture window were not punished, we removed the neural activity after the first lick during the texture window in all following analysis, to avoid any confounding by licking-related activity. As the first lick during choice window triggers outcome, we further aligned each trial to the lick time, and defined the choice window as the 0.5-s period before the first lick in all following analysis. Overall, S1 displayed a higher percentage of texture-responsive neurons compared to other task variables, and this percentage further increased during learning, indicating its role in texture processing. Compared to PPC-RL, PPC-A developed a higher percentage of task-responsive neurons that were distributed across task windows over the learning process (Fig. 2d), indicating its role as a higher-order area in this task. Task-discriminative neurons in these areas showed a similar trend (Fig. 2e). In particular, the percentage of task-discriminative neurons that were jointly responsive to adjacent task windows also increased (Supplemental Fig. 2), potentially benefiting the transition between task phases and allowing for more robust population dynamics during the task. We also investigated the temporal response profile of these task neurons over the course of learning, by comparing the average firing rate of the task-discriminative neurons in naive and expert sessions. As the percentage of task neurons increased during learning, their average response profile also broadened (Supplemental Fig. 2f), suggesting a tiling of task neuron activity during the trials, contributing to longer lasting and more stable population dynamics across S1 and PPC areas. Additionally, the averaged response strength of texture and choice neurons during texture and choice window also increased (Fig. 2f-g). Thus, task-learning was accompanied by an overall recruitment of task neurons, particularly in S1 and PPC-A, as well as extended and enhanced neuronal responses.

### Expanded population response dimensionality and reduced trial variability

Performing a task involves transforming single-neuron activities into a unified sensory representation to guide decisions. During the task, neuronal population activity often resides in a low-dimensional subspace, allowing for robust information representation and reliable behavioral output^12,32–35^. To further understand the changes in the population activity structure regarding variability across neurons and trials, we analyzed the dimensionality of the neuron space and the trial space. For the dimensionality of the neuron space, we performed principal component analysis (PCA) on the population activity with neurons as variables at each time point of the trial, for each session separately (Fig. 3a). Compared to naive sessions, in expert sessions the population dimensionality was increased consistently in S1 and PPC areas in all task windows (Fig. 3b-c). This result suggests that in expert sessions, more neurons with distinct activity patterns are involved during the task. We further analyzed the dimensionality in the trial space by performing PCA on the population activity with trials as variables (Fig. 3d). All three areas, particularly PPC-A, showed decreased dimensionality during the task over the learning process (Fig. 3e-f), indicating decreased trial-to-trial response variability.

**Figure 3.**
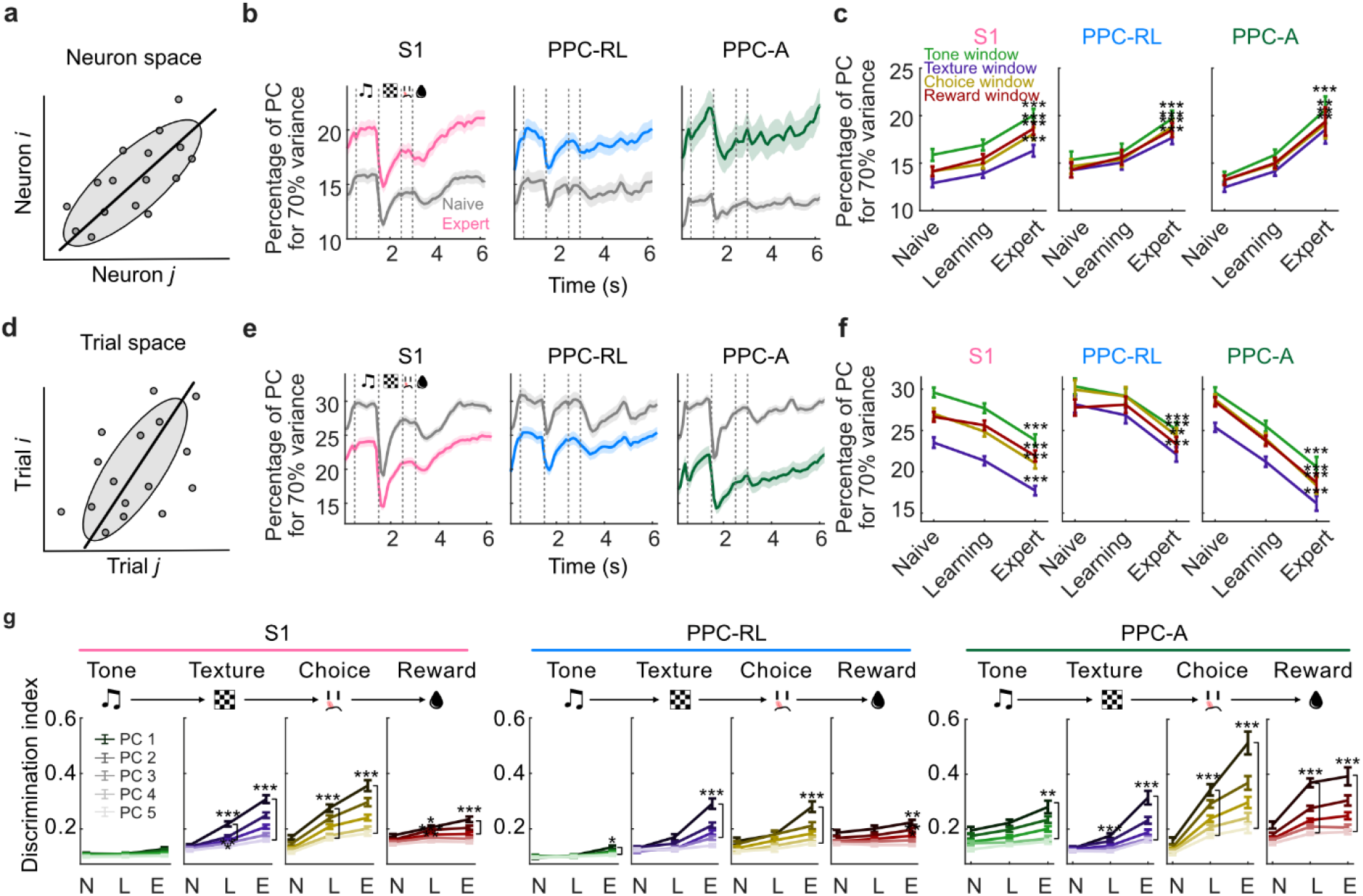
Increased neuronal dimensionality and reduced trial variability during learning. **(a)** Diagram of neuron variance space. PCA was performed with neurons as variables. **(b)** Percentage of principle components (PCs) that explains 70% of variance in neuron subspace, over the trial structure. Dashed lines indicate task windows. **(c)** Quantification of the percentage of PCs in neuron space, during each task window. Colors represent task window. **(d)** Diagram of trial variance space. PCA was performed with trials as variables. **(e)** Percentage of PCs that explains 70% of variance in trial space, over the trial structure. Dashed lines indicate task windows. **(f)** Quantification of the percentage of PCs in trial space, during each task window. Colors represent task window. **(g)** Discrimination index of the top 5 PCs during each task window, regarding the corresponding task variables. Line color saturation represents PC number. (S1: 13 mice, 130, 188, 149 sessions [naïve, N; learning; L, expert, E]; PPC-RL: 13 mice, 70, 77, 104 sessions; PPC-A: 10 mice, 60, 111, 45 sessions; \**p*<0.05, \*\**p*<0.01, \*\*\**p*<0.001; Wilcoxon rank-sum test against naive condition; mean±SEM)

We wondered if the increased neuronal dimensionality is beneficial for carrying task information. To test this, we performed a decoding analysis on the top 5 principal components (PCs) of the neuron space during each task window, using the corresponding task variables (e.g., decoding texture identity during the texture window). All top 5 PCs across S1 and PPC areas showed an increased discrimination index of task variables over the course of learning (Fig. 3g), suggesting that the increased dimensionality could facilitate task information encoding through redundant representation. We conclude that task-learning was accompanied by reduced trial-to-trial variability and enhanced task encoding, potentially through involving more orthogonal representations in S1 and PPC populations.

### Improved task encoding subspaces in S1 and PPC populations during learning

In addition to the global changes in the variance subspace during task-learning, the representation and processing of task information can also be improved through coordinated changes across areas. To examine this aspect, we systematically investigated three additional types of population subspaces during the task: the encoding subspace, the within-area interaction subspace, and the inter-area interaction subspace (Fig. 4a). The encoding subspace captures the optimal encoding direction of task variables; the within-area subspace captures the intrinsic within-area interaction between subsets of neurons in either S1 or PPC; the inter-area subspace captures the interaction between S1 and PPC populations, which we can analyze because we simultaneously imaged from S1 and PPC during the task. We aimed to characterize each type of subspace and directly compare them with each other in the same neuronal space. Since generating within-area subspace requires two subpopulations within an area, we randomly split each population into two subpopulations and computed the three types of subspaces from these subpopulations (Fig. 4a). To reduce the computational complexity, we reduced the subpopulation dimensionality with PCA and kept the top 30 PCs prior to the subspace computation. We repeated this random split procedure 10 times. Because these random splits gave stable results (Supplemental Fig. 3), we averaged the results from all the subpopulations for the following analysis.

**Figure 4.**
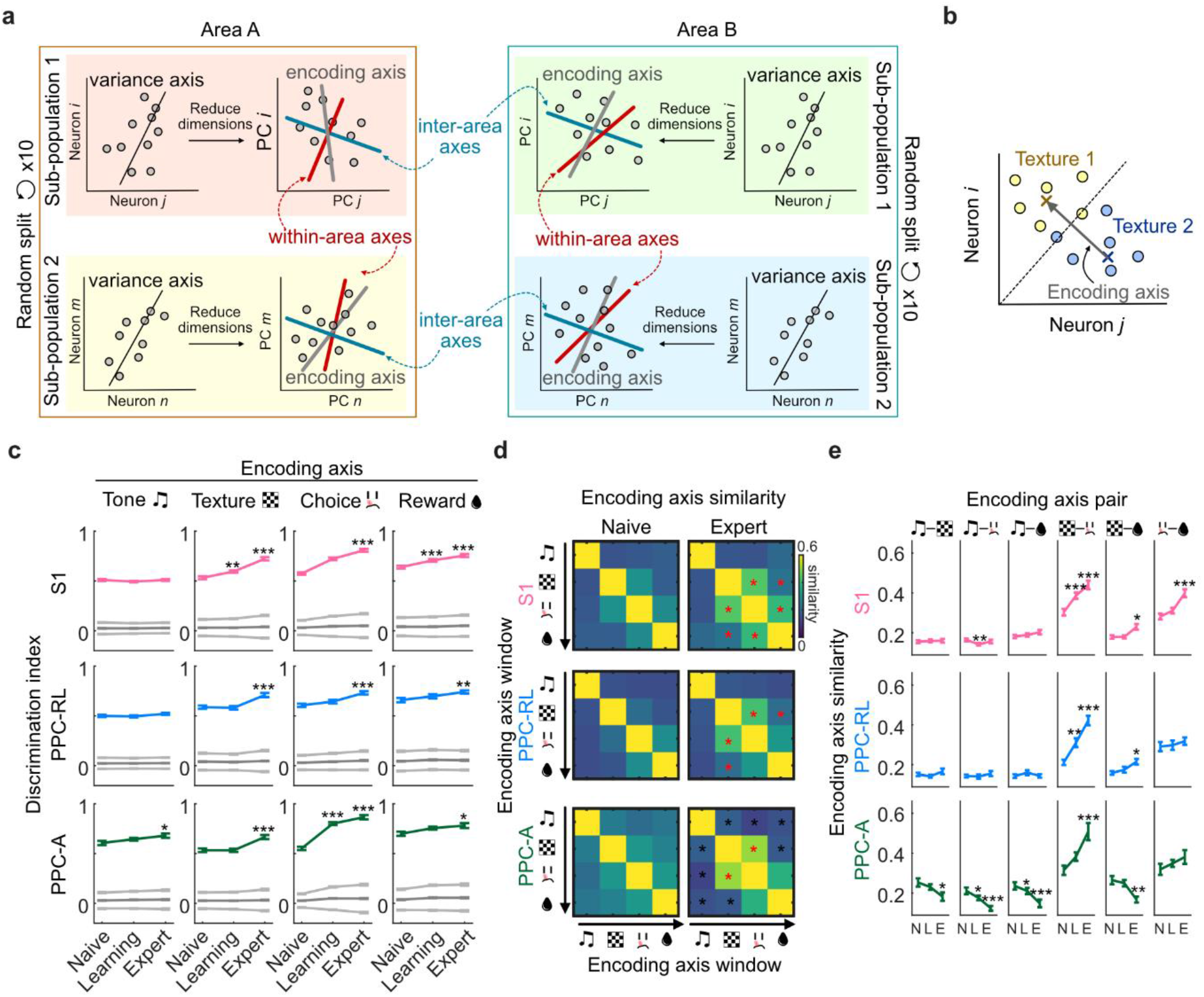
Improved task representation in S1 and PPC encoding subspaces during learning. **(a)** Diagram of generating encoding axis, within-area interaction axis, and inter-area interaction axes in the same reduced 30-dimensional neural space for each subpopulation. **(b)** Example encoding axis scheme for texture axis. Colored dots represent neural activity in texture 1 and texture 2 trials, during the texture window. Crosses represent the means of responses for the two texture types. **(c)** Discrimination index of population activity projection on the encoding axes, for each task variable, in S1, PPC-RL, and PPC-A. **(d)** Color-coded pairwise similarity of encoding axes for all task variable combinations. Symbols on the x- and y-axis indicate task windows. Red and black asterisks indicate that the data from expert condition is significantly higher or lower than the naive condition. **(e)** Quantification of pairwise encoding axes similarity. (S1: 13 mice, 122, 171, 119 sessions [naïve, N; learning, L; expert, E]; PPC-RL: 13 mice, 66, 72, 104 sessions; PPC-A: 10 mice, 56, 99, 33 sessions; C and E: \**p*<0.05, \*\**p*<0.01, \*\*\**p*<0.001; D: \**p*<0.05; Wilcoxon rank-sum test against naive condition; mean±SEM)

We first characterized the encoding subspaces for task variables. We defined the encoding subspace as the axis that captures the difference between the population mean responses to the task variable types^14^, for example, texture 1 vs. texture 2 (Supplemental Fig. 4b). We generated a separate encoding axis for each task variable, using the population activity from the corresponding task window. All encoding axes increased their discrimination index for their target task variable over the course of learning (Fig. 4c), suggesting that the task-related population activity became more separable in this subspace, potentially facilitating the sensory and choice information readout. Furthermore, the encoding axes from adjacent task windows became more aligned over learning (Fig. 4d-e) across S1 and PPC areas, coinciding with the prolonged responses of task neurons as well as the increase in jointly responsive task neurons (Fig. 2f-g; Supplemental Fig. 2). In PPC-A, the encoding subspace from non-adjacent task windows even became more distinct from each other. These results suggest that the encoding subspace was reorganized over the learning process to optimize the representation of task information.

### Improved alignment of encoding subspaces with within-area interaction subspaces

We next characterized the within-area interaction subspaces. To do so, we applied canonical correlation analysis (CCA) to the two subpopulations from each area (Fig. 5a). CCA is a statistical method for quantifying the relationship between two sets of variables, and has been used previously to study the interaction between neuronal populations^23,36–38^. Conceptually, CCA finds pairs of projection axes from the two populations that maximize the correlation between the projections. Similar to PCA, CCA finds orthogonal sets of projection axes within each population. These axes are ordered with descending correlation values between the paired projections, along the canonical dimensions. The top pair of projection axes thus represents the optimal interaction subspace between the two populations, capturing the strongest canonical correlation.

**Figure 5.**
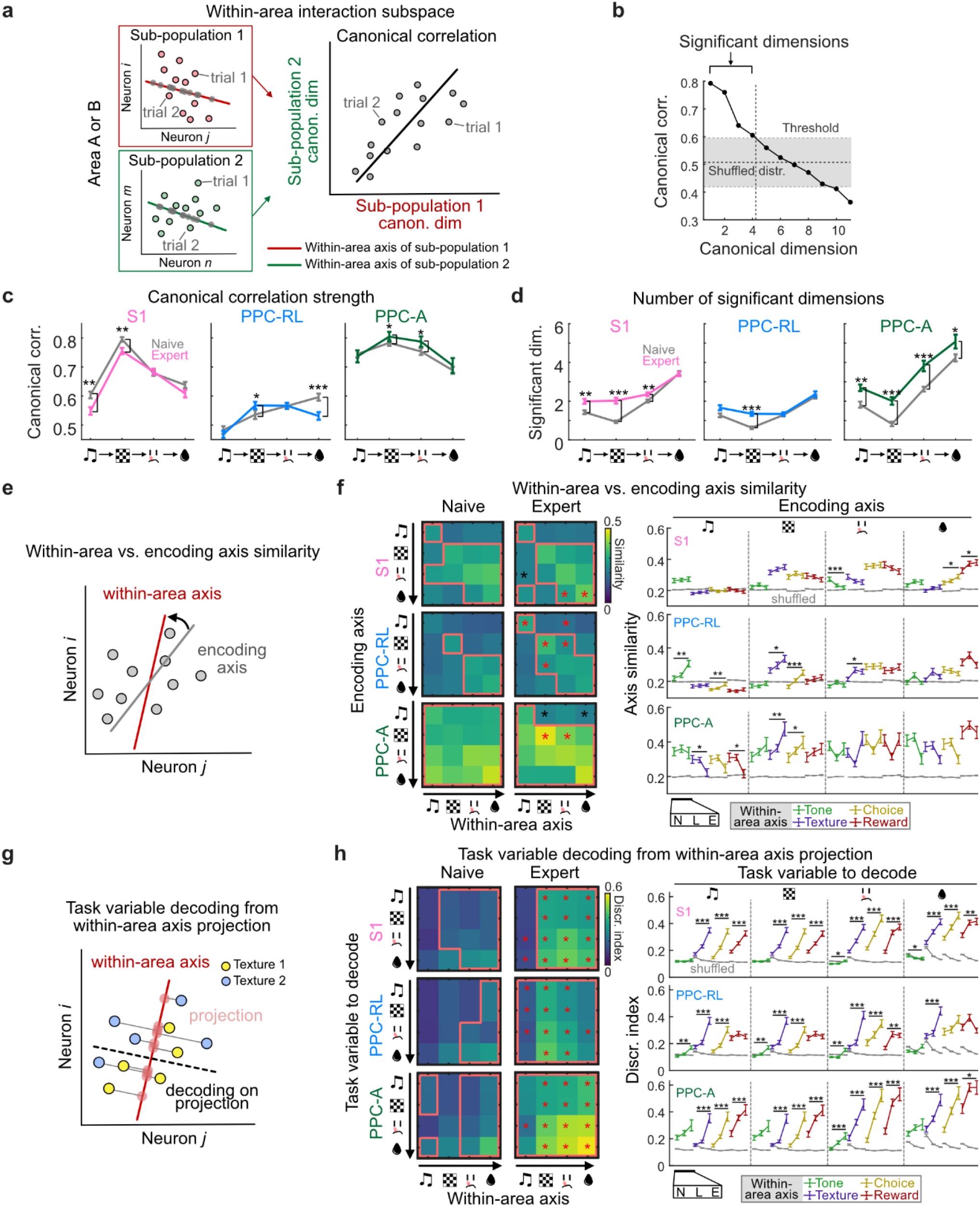
Task encoding axes aligned with PPC within-area axes during learning. **(a)** Diagram of using CCA to generate pair of within-area interaction axes from sub-populations of the same area, which maximizes the canonical correlation between the projected data from each sub-population. **(b)** Example of identifying significant CCA dimensions. Black line represents canonical correlation in descending dimensions; gray zone represents confidence interval from shuffled models. **(c)** Canonical correlation strength in naive and expert condition for each task window, in S1, PPC-RL, and PPC-A. **(d)** Number of significant dimensions in naive and expert condition for each task window, in S1, PPC-RL, and PPC-A. **(e)** Diagram of comparing within-area interaction axis with encoding axis, using cosine similarity. **(f)** Left: pairwise similarity between within-area interaction axis (x-axis) and encoding axis (y-axis), across task windows. Red boxes indicate that the data is above the corresponding shuffled distribution; red and black asterisks indicate that the data from expert condition is significantly higher or lower than the naive condition. Right: quantification of axis similarity. Line colors represent the task window of the within-area axis; gray lines represent 95% quantile of shuffled data. **(g)** Diagram of computing decoding performance from population activity projection onto within-area interaction axis. This example shows texture decoding, using neural activity during the texture window to project onto the within-area axis. **(h)** Left: discrimination index of task variables (y-axis), using projection onto within-area axis from each task window (x-axis). Lines and asterisks represented as in **f**. Right: quantification of discrimination index. Colored lines represent the task window of within-area axis. (S1: 13 mice, 122, 171, 119 sessions [naive, learning, expert]; PPC-RL: 13 mice, 66, 72, 104 sessions; PPC-A: 10 mice, 56, 99, 33 sessions; \**p*<0.05, \*\**p*<0.01, \*\*\**p*<0.001; **f** and **h** left panels: \**p*<0.05; Wilcoxon rank-sum test against naive condition; mean±SEM)

Given the two subpopulations from the same area, their top canonical correlation value represents the within-area interaction strength, as it reflects the covariance between the subpopulations. Among the three areas, PPC-A overall displayed the highest correlation strength (Fig. 5c). Compared to naive sessions, the within-area interaction strength slightly decreased in expert sessions in S1, but slightly increased in PPC, particularly in PPC-A (Fig. 5c). We further investigated the within-area subspace dimensionality by identifying the number of significant interaction dimensions, defined as the number of canonical dimensions with a correlation value higher than the shuffled threshold (Fig. 5b). The shuffled distribution was generated by taking the top canonical correlation from shuffled models, where the trial correspondence was randomized. Among the three areas, PPC-A had the highest within-area subspace dimensionality, in line with it being relevant as a higher-order area in this task. Over the course of learning, the within-area subspaces in S1 and PPC-A significantly increased their dimensionality across task windows, while PPC-RL showed an increase only during the texture window (Fig. 5d). We conclude that task-learning expanded and enhanced within-area interactions, particularly in PPC-A.

We then wondered whether the within-area interaction axis contains more task-relevant information over the learning process, through improved alignment with the task encoding axis. To answer this question, we computed the cosine similarity between the encoding axis and the top within-area axis computed from the same subpopulation (Fig. 5e). We performed this comparison for each pair of axes from all task windows. In S1, the alignment between the encoding axes and the within-area axes from texture and choice windows remained stable over the course of learning. In PPC areas, particularly in PPC-A, the texture encoding axis became more aligned with the within-area axes during the texture and choice windows (Fig. 5f). To understand whether this improved alignment was due to changes in the within-area subspace structure, we also computed the similarity between pairs of within-area axes over the trial time. We found that the within-area axis structure remained stable in PPC-A, and even became more distinct in S1 and PPC-RL over the learning process (Supplemental Fig. 4a). Considering that the encoding axis structure was reorganized and optimized during learning (Fig. 4d-e), these results suggest that PPC task encoding subspaces reconfigured to align with their within-area subspaces during learning, potentially contributing to improved task representation by the local populations.

To further test whether the within-area subspace developed more task information over learning, we projected the population activity onto the within-area axes and performed a decoding analysis for each task variable (Fig. 5g). In expert sessions, the discrimination index of within-area activity was improved across S1 and PPC areas, particularly in PPC-A (Fig. 5h), in agreement with the increased number of task neurons as well as improved population encoding performance (Fig. 2-3). Therefore, task-relevant information was enhanced across within-area subspaces in S1 and PPC during learning.

### Improved alignment of encoding subspaces with inter-area interaction subspaces

Our experimental design of simultaneous imaging from S1 and PPC areas allowed us to also probe the inter-area interaction subspaces between them. To do so, we applied CCA to the subpopulations from S1 and PPC-RL, or from S1 and PPC-A (Fig. 6a). As for the within-area subspace analysis, we computed the top canonical correlation strengths and the number of significant dimensions of interactions. Over the course of learning, the interaction strength between S1 and PPC-RL slightly decreased, while their interaction subspace dimension expanded during texture window. In contrast, S1 and PPC-A interaction strength remained stable and consistently higher compared to S1 and PPC-RL interactions. Additionally, the interaction subspace dimensionality increased for S1 and PPC-A during tone, texture and choice windows (Fig. 6b-c), suggesting a specific co-involvement of S1 and PPC-A in the task. We then compared the alignment between inter-area axes and encoding axes for each pair of areas, measured by the cosine similarity (Fig. 6d). Both PPC-RL and PPC-A showed improved alignment of encoding axes and inter-area axes with S1 over the learning process. In PPC-RL, the texture and choice encoding axes became more aligned with the inter-area axes with S1 after the texture onset (Fig. 6f). In PPC-A, this improvement in alignment occurred earlier during the trial structure, before the texture window, where the inter-area axes during tone window developed texture and choice information over learning (Fig. 6g). To understand whether these changes in alignment can be explained by the changes of the intrinsic inter-area axis structure, we also analyzed the similarity between pairs of inter-area axes across task windows. We found the inter-area interaction structure between S1 and PPC areas mostly remained stable over the learning process (Supplemental Fig. 4c-d). Considering the learning-related reorganization in the encoding subspace structure (Fig. 4d-e), these results suggest that the task encoding subspaces became better aligned with S1-PPC interaction subspaces.

**Figure 6.**
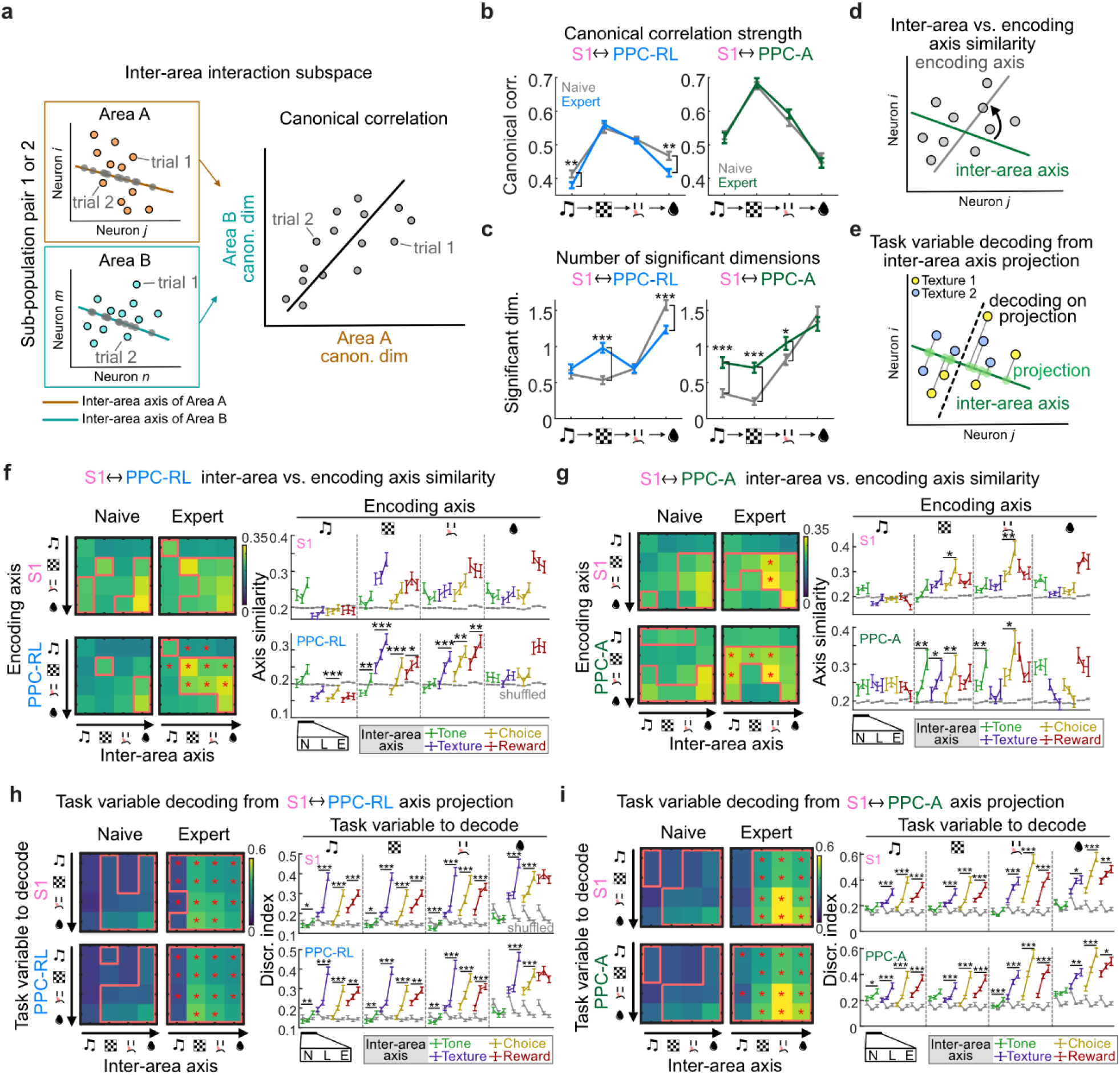
Task encoding axes aligned with S1 and PPC inter-area axes during learning. **(a)** Diagram of using CCA to generate pair of inter-area interaction axes from sub-populations of two different areas (Area A and B), which maximizes the canonical correlation between the projected data from each sub-population. **(b)** Canonical correlation strength in naive and expert conditions for each task window, for S1 and PPC-RL interaction (left) and S1 and PPC-A interaction (right). **(c)** Number of significant dimensions in naive and expert conditions for each task window. **(d)** Diagram of comparing inter-area axis with encoding axis. **(e)** Diagram of computing decoding performance from population activity projection onto inter-area axis. This example shows texture decoding, using neural activity during the texture window to project onto within-area axis. **(f)** Inter-area axis similarity with encoding axis in S1 (top) and PPC-RL (bottom). Left: pairwise similarity between inter-area axis (x-axis) and encoding axis (y-axis), across task windows. Red boxes indicate that the data is above the corresponding shuffled distribution; red or black asterisks indicate that the data from expert condition is significantly higher or lower than the naive condition. Right: quantification of similarity. Line colors represent inter-area axis windows; gray lines represent 95% quantile of shuffled data. **(g)** Same plots as in **f** for inter-area axis similarity with encoding axis for S1 (top) and PPC-A (bottom). **(h)** Task variable decoding from S1 and PPC-RL inter-area axis projection. Left: discrimination index of task variables (y-axis), using projection onto inter-area axis from each task window (x-axis). Right: quantification of discrimination index. Colored lines represent inter-area axis windows. **(i)** Same plots as in **h** for task variable decoding from S1 and PPC-A inter-area axis projection. (S1 and PPC-RL interaction: 13 mice, 66, 72, 86 sessions [naive, learning, expert]; S1 and PPC-A interaction: 10 mice, 56, 99, 33 sessions; \**p*<0.05, \*\**p*<0.01, \*\*\**p*<0.001; **f, g, h, i** left panels: \**p*<0.05; Wilcoxon rank-sum test against naive condition; mean±SEM)

We further tested whether the inter-area subspace developed more task information over learning, following its improved alignment with task encoding subspaces. As above, we projected the population activity onto the inter-area axes and performed decoding analysis for each task variable (Fig. 6e). We found that the projected inter-area activity of S1 and PPC showed improved discriminability of task variables, particularly for S1 and PPC-A interaction (Fig. 6h-i). Therefore, the interaction between S1 and PPC areas carried enhanced task-relevant information over learning.

### Reduced population dimensionality and misaligned subspaces during mistakes

During the expert phase, mice still made mistakes (Fig. 1c). In these expert incorrect trials, the licking pattern as well as the pupil and face movement of mice were different from the incorrect trials in naive sessions, but were overall similar with the expert correct trials despite slight differences (Fig. 1g-i; Supplemental Fig. 5). We wondered whether these expert incorrect choices can be explained by changes in the underlying task representation on both single-neuron and population levels. We first compared the activity of task neurons during correct and incorrect trials in expert sessions. Compared to correct trials, the response of task-discriminative neurons showed reduced activity strength as well as shortened response profile in incorrect trials (Fig. 7a-b), suggesting a less stable neural code. While the population dimensionality in the neuron space and in the trial subspace was minimally affected by incorrect behaviors during the expert phase (Supplemental Fig. 6a-d), the encoding subspace of S1 and PPC populations in incorrect trials showed reduced task variable discriminability (Fig. 7c). Consistently, the top PCs in neuron space also carried less task information on average during incorrect trials across S1 and PPC areas (Supplemental Fig. 6e).

**Figure 7.**
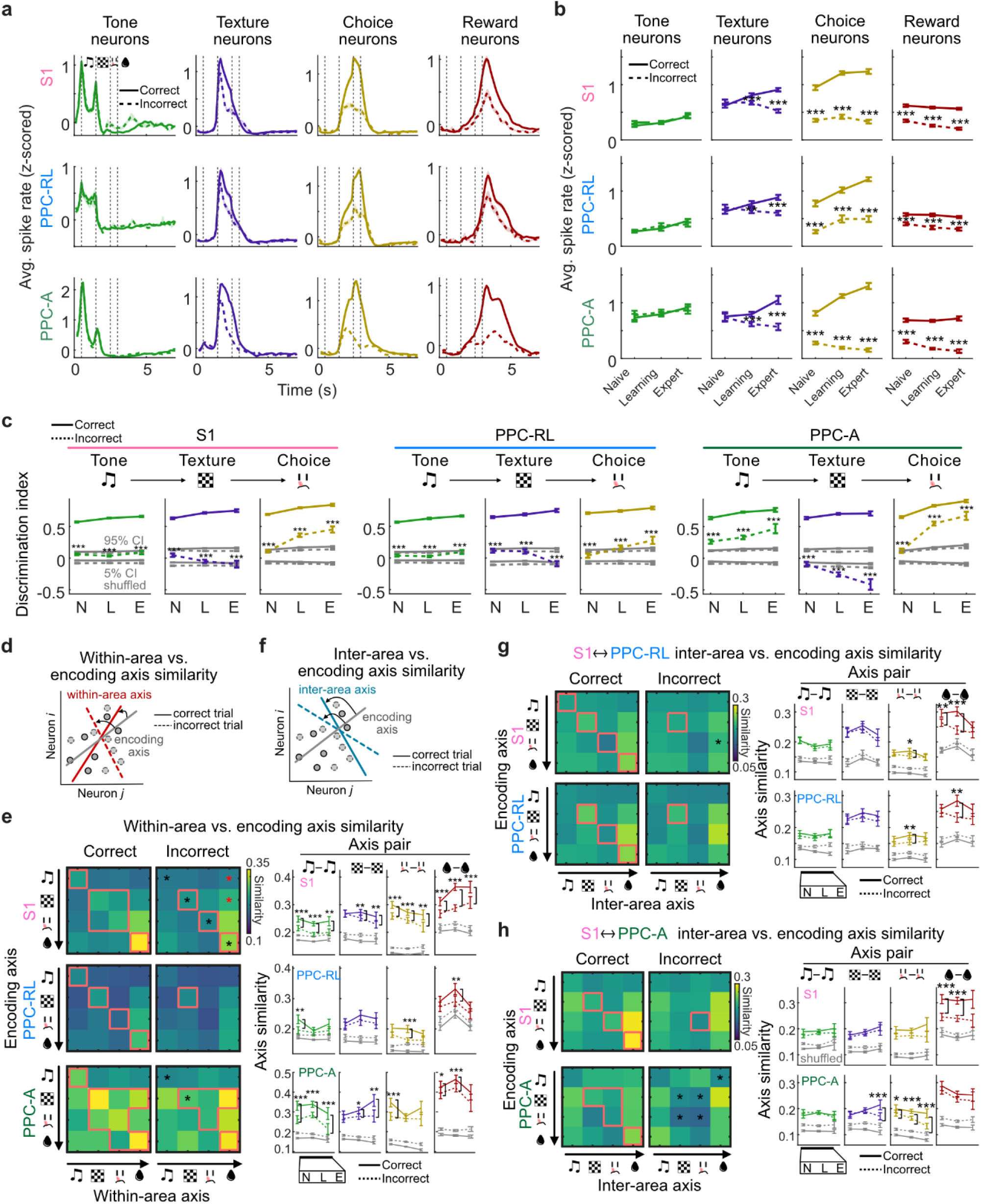
Disrupted task-related neural representations during incorrect trials. **(a)** Session-averaged spike rates of task-discriminative neurons over trial time, during correct (solid lines) and incorrect (dashed lines) trials. Spike rates were z-scored within each session. Gray lines represent naive condition, colored lines represent expert condition. Dashed vertical lines indicate task windows. **(b)** Quantification of average spike rates of task-discriminative neurons during correct (solid lines) and incorrect (dashed lines) trials, in different task windows. Colors represent the quantified task window. **(c)** Discrimination index of population activity projections on the encoding axes during correct (solid lines) and incorrect (dashed lines) trials, for each task variable, in S1, PPC-RL and PPC-A. Gray lines represent shuffled data confidence interval (CI). **(d)** Diagram of comparing within-area axis with encoding axis, in correct and incorrect trials. **(e)** Within-area axis similarity with encoding axis in expert condition. Left: pairwise similarity between within-area axis (x-axis) and encoding axis (y-axis), in correct (left panels) and incorrect (right panels) trials. Red lines indicate that the data is above the corresponding shuffled distribution; red or black asterisks indicate that the data from expert condition is significantly higher or lower than the naive condition. Right: quantification of similarity. Solid lines represent correct trials, dashed lines represent incorrect trials, gray lines represent shuffled data. **(f)** Diagram of comparing inter-area axis with encoding axis, in correct and incorrect trials. **(g)** Inter-area axis similarity with encoding axis in expert condition for S1 (top) and PPC-RL (bottom). Left: pairwise similarity between inter-area axis (x-axis) and encoding axis (y-axis), in correct (left panels) and incorrect (right panels) trials. Right: quantification of similarity. **(h)** Same plots as in **g** for inter-area axis similarity with encoding axis for S1 (top) and PPC-A (bottom). (S1: 13 mice, 149, 156, 48 sessions [naive, learning, expert]; PPC-RL: 13 mice, 79, 63, 33 sessions; PPC-A: 10 mice, 70, 93, 15 sessions; S1 and PPC-RL interaction: 13 mice, 79, 63, 33 sessions; S1 and PPC-A interaction: 10 mice, 70, 93, 15 sessions; \**p*<0.05, \*\**p*<0.01, \*\*\**p*<0.001; **e, g, h** left panels: \**p*<0.05; Wilcoxon signed rank paired test between correct and incorrect trials; mean±SEM).

Finally, we investigated the changes in within-area and inter-area subspaces during incorrect trials. We generated within-area and inter-area subspaces for correct and incorrect trials separately, using matching number of trials (Methods). While both within-area and inter-area interaction strength and dimensionality were minimally affected by incorrect behaviors (Supplemental Fig. 6f-k), these axes were less aligned with the task-encoding axes compared to correct trials (Fig. 7d-h), particularly in S1 and PPC-A. In addition, population activity projected onto both within-area and inter-area axes carried less task information (Supplemental Fig. 7), particularly for the inter-area axes of S1 and PPC-A. Together, these results suggest that behavioral mistakes in expert mice were associated with less reliable neuronal responses and misaligned encoding and intrinsic interaction subspaces, potentially leading to erroneous task information processing and behavioral output.

## Discussion

Using simultaneous multi-area two-photon imaging combined with a sensory discrimination task, we showed that task-learning was associated with multiple changes in S1 and the higher association areas PPC-A and PPC-RL, both of which were required for optimally performing the task. On the single-neuron level, additional task neurons were recruited, and the task-related neuronal responses were enhanced. On the population level, the response dimensionality was increased, and task representations were enhanced in the corresponding orthogonal subspaces. The population readout of task information was also enhanced through reconfigured and optimized task encoding axes. Furthermore, learning this task preferentially involved PPC-A throughout the trial structure, opening additional interaction subspaces within PPC-A and between S1 and PPC-A, while PPC-RL was involved mostly in texture processing. Over the course of learning, task encoding subspaces became better aligned with the within- and inter-area subspaces in PPC areas, improving information readout in these channels. Incorrect choices of expert mice were accompanied by less reliable neuronal responses and misaligned encoding and intrinsic interaction subspaces, potentially contributing to the behavioral mistakes.

While both dimensionality increase and decrease of neuronal population response have been observed during learning^32,39–41^, studies have also shown that storing task information in orthogonal subspaces can improve the overall task representations by the population^10,13,42^. Using these separate subspaces not only increases the stimulus encoding reliability, but also allows separate task information, or even separate tasks, to be stored together^10,13,42^. In our study, we observed an increase in the neuronal dimensionality, as well as an increase in both sensory and choice information in these subspaces, corresponding to an overall improvement of task representation. On the other hand, reduced task information encoded by fewer subspaces was linked with incorrect choices. This suggests that learning this sensory discrimination task benefited from higher population dimensionality and redundant information within S1 and PPC.

At the local population level, task-learning can be accompanied by enhanced task information readout in both primary sensory areas and higher areas^11,12,14,43^. This can be achieved through enhancing neural coding consistency^44^ and re-aligning the readout dimensions with the more stable intrinsic within-area subspaces^11^. Enhancing the information readout in a higher area, without changing the primary area responses, is sometimes sufficient to improve task performance^43^. In our study, we found evidence for improved task information in both S1 and PPC. As the within-area subspaces in PPC strengthened and expanded, the task encoding dimensions also better aligned with the PPC within-area subspaces, coinciding with an improved task representation, particularly in PPC-A. This supports both mechanisms of enhanced coding consistency and optimized population readout through re-alignment.

Another possible mechanism of learning is through enhanced signal propagation between areas. This can be achieved through stronger shared signals between areas and by better aligning the task information and interarea communication subspaces^20,45^. For example, studies have shown that task-learning increases shared dimensionality among motor cortex neurons^20^ and that top-down signals can change the population interaction structure dynamically during a behavioral task^21^. However, although the inter-area communication subspaces can be flexible^46,47^, task-learning does not require reconfiguration of these subspaces. In many cases, the intrinsic within- and across-population activity structures remain stable and impose constraints on task learning^16,22^. Improving task representation along the inter-area subspaces without changing them, e.g., by regulating the internal states such as attention, is sufficient to improve task performance^17,45^. Our results support this mechanism by showing that task encoding subspaces gradually aligned with the S1 and PPC inter-area subspaces without changing the interaction subspace structure.

Task-learning often involves changes in specific pathways^6,18,19,48^. Our study is focused on learning-related changes in S1 and PPC areas. While both PPC-RL and PPC-A were engaged in this task and required for optimal task performance, we found that the S1-to-PPC-A pathway was more involved than the S1-to-PPC-RL pathway during learning, particularly during the earlier tone window. As a higher association area, the PPC is important for routing sensory information in behavioral tasks, with its activity emerging during learning as an intermediate step of sensory information flow^26,28^. Our study adds to the accumulating evidence that different PPC subregions have distinct roles^23,26^. The higher engagement of PPC-A than PPC-RL area in our study is possibly due to the task design, where a specific tone was paired with a fixed subsequent texture, allowing mice to form specific tone-texture associations^23^. However, PPC is a highly flexible area, and a change in the task design could strongly influence whether and how PPC is involved^49^.

In our specific task design, there are two types of learning: sensory association learning, which links tone stimuli to the paired subsequent texture stimuli, and sensorimotor learning, where the tone-texture sequence is associated with a specific choice. We observed some differences regarding these two types of learning. During the tone window, which represents sensory association learning, the populations showed less prominent increases in discrimination ability and number of task-responsive neurons, but more noticeable increases in the population dimensionality as well as the S1 to PPC-A interaction dimensions. During the texture window, which is prior to the choice window and thus represents sensorimotor learning, the population showed increased percentage of task neurons and improved discrimination ability, as well as increased within- and inter-area dimensions. These results suggest that distinct mechanisms exist for different types of learning, which will require further studies to fully understand these processes.

Overall, our results show that task-learning is achieved through coordinated changes on multiple levels, across neocortical areas. These changes include local recruitment of task neurons, and the improved alignment of task-relevant information with the intrinsic interaction subspaces within and across areas, resulting in improved sensory processing and refined behavioral output. These findings highlight the complexity of learning processes, and prompt future research to further understand the specific mechanisms and their impact on neural computation and behavior underlying learning.

## Acknowledgements

We thank Philipp Bethge for managing the transgenic mouse lines, Fabian Voigt and Hansjörg Kasper for help with optics, and Martin Wieckhorst for the behavior training software. We also thank Matthias Tsai and Christopher Lewis for their feedback on the manuscript. This work was supported by a Sinergia grant from the Swiss National Science Foundation (CRSII5_180316; to F.H.) and a UZH Forschungskredit grant (K-41220-07-01; to S.H.). S.H. received an Ambizione Grant from the Swiss National Science Foundation (PZ00P3_216312). F.H. received funding from the University Research Priority Program (URPP) “Adaptive Brain Circuits in Development and Learning” (AdaBD).

## Author contributions

S.H. and F.H. conceived the study and designed the experiments. S.H. performed the experiments and analyzed the data. S.H. and F.H. wrote the manuscript.

## Methods

All procedures of animal experimentation were carried out according to the guidelines of the Veterinary Office of Switzerland and following approval by the Cantonal Veterinary Office in Zurich (licenses 234/2018, 211/2018, 141/2022).

### Mice and dataset

Part of the dataset and method has been previously published^23^. Mice were housed on a 12-h reversed light/dark cycle at an ambient temperature of between 21°C and 23°C, with humidity level between 55% and 60%. A total of 16 mice were included in this study. Mice belonged to one of the following transgenic strains: RasGRF2a-dCre;CamK2a-tTA;TITL-GCaMP6f (M10, M11, M12, M25, M26, M28, M29, M33, M34, M35), GP5.17(C57BL/6J-Tg(Thy1-GCaMP6f)GP5.17Dkim/J,

Jackson Laboratory 025393) (M14, M15), Snap25-IRES2-Cre-D;CamK2a-tTA;TITL-GCaMP6f (M17), RasGRF2a-dCre;tTA2-GCaMP6f (M30, M38, M40). All mice expressed GCaMP6f in layer 2/3 pyramidal neurons of the neocortex. Both sexes were included in this study (male: M10, M14, M15, M25, M26, M30, M33, M34, M35; female: M11, M12, M13, M17, M28, M29, M38, M40). Among the 16 mice, S1-PPCA imaging was performed on 9 mice (M25, M26, M28, M29, M30, M33, M34, M35, M38, M40), with 5 mice imaged since naive phase (M33, M34, M35, M38, M40), the rest imaged during late learning and/or expert phase; S1-PPCRL imaging was performed on 14 mice (M10, M11, M12, M14, M17, M25, M26, M28, M29, M30, M33, M34, M35, M40), with 9 mice imaged since naive phase (M10, M11, M12, M14, M17, M25, M26, M28, M29), the rest imaged during late learning and/or expert phase. Mice were 2.5-4 months old at the beginning of behavior training, and 3-5 months old at the time of imaging.

### Surgical procedures

We performed a craniotomy over S1 and PPC in the left hemisphere of all mice. During surgery, mice were anesthetized with 2% isoflurane mixed with oxygen and maintained at 37°C body temperature. Mice were treated with analgesia medication (Metacam, 5 mg/kg, s.c.; lidocaine gel over the skull skin) before exposing the skull. Then, a 4-mm round cranial window was made and covered with a glass coverslip using dental cement (Tetric EvoFlow). A light-weighted head-post was mounted on the skull using dental cement. Mice were continually monitored after surgery for at least three days. For strains expressing destabilized Cre (dCre), we induced GCaMP6f expression by intraperitoneally administering trimethoprim (TMP, Sigma T7883; in Dimethyl sulfoxide (DMSO, Sigma 34869) at 100 mg/ml; 150 mg TMP/g body weight) at least one week before imaging started.

### Behavior training

Mice recovered for at least 1 week before the behavior training started. During the first phase of training, mice were handled by the experimenter for several days until showing no sign of stress, then they were gradually accustomed to head fixation. Next, mice were put on water scheduling, and they were introduced to the behavior setup. During the first 2-3 sessions, mice obtained sugar water auto-reward after the choice tone (2 beeps at 3 kHz of 50-ms duration with 50-ms interval), from one of the two lick ports. Once they learned to lick after the choice tone, we introduced the full task.

Behavior training was conducted using custom LabView software. Each trial started with either a low tone or a high tone (10 kHz or 18 kHz, 6 repetitions, 50-ms duration and 50-ms intervals), followed by a matched smooth or a rough texture (sandpaper, P100 vs. P1200 for M10-17, P280 vs. P800 for M25-40) carried by a rotary motor, mounted on a linear stage. Both the tone and the texture were presented for 1 second. Then, the choice window started with the choice tone described above, lasting for up to 2 seconds. As soon as mice licked during the choice window, it was terminated and the reward window started. Each correct choice led to a small sugar water reward (∼4 µl) on the respective water spout; incorrect choices were neither rewarded nor punished. The inter-trial interval was randomly distributed between 4-8 seconds.

Training started with 2-3 sessions of auto-reward, where the reward was automatically delivered from the correct lick port, no matter what the mice chose. Once the mice were accustomed to the task structure, formal training started. During training, we adopted a “repeat incorrect” strategy, where incorrect trials were followed by the same tone-texture stimulus pair until the mouse chose correctly. To motivate the mice, approximately 10% of miss trials were auto-rewarded after the choice. Each day, the training lasted as long as the mouse was actively engaged in the task, typically 100-500 trials. Mice were trained once per day for 5-6 days a week. Weight, health, and water intake were monitored daily. All training was performed in the dark, with mice continuously monitored with a CMOS infrared-sensitive camera (Basler acA1440-220um) under 850-nm infrared LED background illumination. Behavioral videos were recorded at 50 Hz, and body movement was computed using frame-to-frame correlation. The pupil was constrained by a small UV LED (385 nm, Thorlabs LED385L) positioned close to the eye, and illuminated by the two-photon near-infrared laser. We tracked the pupil diameter by binarizing the pupil image and fitting an ellipse to the pupil region. Body movement and pupil diameter were both smoothed with a median filter of 200-ms width.

To account for the differences in trial number for each day during learning, we split the training days into sessions of 80-120 trials (median: 120 trials; mean ± SD: 116 ± 9 trials) and computed all analysis based on sessions. During the expert phase, mice went through mismatch experiments, where we studied predictive processing, as previously published^23^. Briefly, 10-30% percent of the trials were mismatch trials where tone-texture pairing was inverted. Rewards were delivered according to the texture identity, which led to minimal confusion for mice. Since the mismatch trials were infrequent, and we implemented the repeat incorrect strategy in these experiments to reinforce normal trial pairing, we also included these datasets for analysis, but excluded all mismatch trials as well as sessions with lower performance (<70%).

### Optogenetics

For the optogenetics experiment, we trained 8 VGAT-ChR2-EYFP mice to perform the behavioral task. Before behavior training, we implanted optical fibers (400-µm diameter, NA 0.39) bilaterally above PPC-A and PPC-RL, through a small cranial window. The coordinates used to determine PPC-A and PPC-RL are (-1.8, 2.25) and (-2.5, 3.35), respectively (from Bregma and midline, in mm). Laser light (470 nm, 1 mW above cortex; Thorlabs M470F4) was modulated by a 40-Hz square wave (50% duty cycle) and delivered throughout the texture window for 1 second. In each session, either bilateral PPC-A or bilateral PPC-RL were inhibited in 50% of the trials. Due to the performance fluctuation of this transgenic strain, we excluded the non-expert subsessions during analysis. At the end of all optogenetics sessions, we conducted tone-only and texture-only sessions, where either the tone or the texture were omitted while maintaining the original temporal structure of trials. Optogenetics was performed during the texture window, as described above.

### Sensory mapping

The exact locations of S1, PPC-A, and PPC-RL were determined using widefield sensory mapping and retinotopic mapping, as previously reported^23^. Briefly, for widefield mapping, mice were lightly anesthetized and presented with visual, whisker, and hindlimb stimuli (30 repetitions for each modality) on the contralateral side to the imaging window, while we performed simultaneous widefield imaging. In the widefield imaging system, we used a blue LED light source (Thorlabs; M470L3) and an excitation filter (480/40 nm BrightLine HC), a 4x objective (Thorlabs TL4X-SAP, NA 0.2) for imaging, an emission filter (529/24 nm, BrightLine HC), and a CMOS camera (Hamamatsu Orca Flash 4.0) for collection. For retinotopic mapping, a drifting spherically-corrected checkerboard visual stimulus of four cardinal directions (10 repetitions each direction) was presented on an LED screen (Adafruit Qualia 9.7” DisplayPort Monitor, 2048×1536 pixel resolution) across the visual field of the mice. The retinotopic map was calculated using a previously reported analysis pipeline^50^. The final locations of S1, PPC-A and PPC-RL were determined by optimally aligning the sensory map and retinotopic map together to the Allen Mouse Common Coordinate^51^.

### Two-area two-photon imaging

Two-area two-photon imaging was performed using a previously reported custom-built microscope^24^. The simultaneous two-area imaging was achieved through a temporal multiplexing scheme, where the laser pulses from a Ti:sapphire laser (Mai Tai HP DeepSee, Spectra-Physics) was split in two temporally interleaved copies, each directed through an independently movable unit to a separate field of view. Two-photon imaging was performed at 920-nm excitation with a green emission filter (510/42 nm bandpass), through a 16x objective (N16XLWD, Nikon, NA 0.8). In each area, we performed volumetric imaging using an electrically tunable lens (Optotune EL-10-30-C), from 3 different depths in layer 2/3, separated by 40-50 µm, with typically 450×500 µm FOV size, at a resolution of typically 370×256 pixels. For two areas and three imaging depths per area, the volume rate was typically 9.3 Hz. During learning, we followed the same FOVs in each mouse, guided by the blood vessels and landmark neurons, with minor shifts each day. In expert experiments, we slightly changed the depths in each session to cover slightly different populations. Due to the dense labeling of the GCaMP6f transgenic mice, the long learning curve, and the shifts in FOVs across days, we could not identify enough neurons throughout the whole learning curve, therefore we did not seek to match the neurons across days.

### Processing of two-photon imaging data

We used Suite2p^31^ to perform rigid motion correction on the raw data, model-based background subtraction, neuron identification, fluorescence extraction, and neuron classification. This pipeline outputs the raw fluorescence traces, the neuropil traces, and the deconvolved spike rates of identified neurons. We manually curated each imaging session and discarded non-neuronal structures or low-quality ROIs. All analysis was performed using the z-scored deconvolved spike rates, where the spike rate of each neuron was normalized to have zero mean and unit standard deviation.

Due to fluorescence signal bleed-through between two areas or between two adjacent imaging depths within each area, we carefully removed potentially redundant neurons that were highly correlated with neighboring neurons. We identified potential redundant neuron pairs that fulfilled all the below criteria: (1) spike rate correlation above 0.5; (2) lateral distance between centroids below 5 µm regardless of depths; (3) appeared in adjacent imaging depths in the same imaging area (signal bleed-through in the same area from adjacent imaging planes), or appeared in the same imaging depths in both imaging areas (signal bleed-through across areas from the same imaging plane). In these duplicated neuron pairs, we kept the neuron with highest average fluorescence level, and discarded the one with less fluorescence. In each imaging session, the number of redundant neuron pairs were typically below 5.

Due to the variable length of the choice window, we defined choice window as the 0.5-s time period before the lick event that triggered the reward window. To exclude lick-related activity during the texture window, we identified all early licks during the texture window, and discarded the subsequent activity in these trials until the choice window. We performed this procedure in all analyses except for the sliding window PCA analysis, where discarding early licks during the texture window introduced artificial population dimensionality shrinkage.

### Task neuron analysis

We identified task-tuned neurons by testing the activity level of individual neurons across task windows against a null distribution. We first denoised the spike rates by a small Gaussian kernel (3 frames, sigma=1); then, for neuron *N*i during task window *T*j, we compared its average activity within the window against a null distribution of average activity level, generated by randomly sampling the same number of frames outside of window *T*j for 100 times. The neuron *N*i is identified as responsive in task window *T*j if its activity in *T*j was significantly higher than the null distribution (one-tailed Wilcoxon Rank Sum test; p<0.05).

To identify task-discriminative neurons, we tested the activity of responsive neurons during different values of each task variables (tone, texture, choice, and reward). For each task variable (for example texture), there are two possible values *s*1 and *s*2 (texture 1 and texture 2). For neuron *N*i, we compared its average activity within the corresponding task window (texture window in this example) between *s*1 trials and *s*2 trials. Neuron *N*i was identified as a discriminative neuron if its activity was significantly higher (Wilcoxon Rank Sum test, p<0.05) in *s*1 or *s*2 trials. We performed this procedure for all four task variables in the corresponding task windows.

### Discrimination analysis

To calculate the discrimination ability of principal components (PCs) or axis projections, we calculated a standard receiver operating characteristics (ROC) curve using each task variable and computed its area under the curve (AUC). The discrimination index was defined as DI=(AUC-0.5)×2.

### Encoding axis analysis

We aimed to compare the encoding axis, the within-area interaction axis, and the inter-area interaction axis within the same space. Since the within-area axis requires two subpopulations from the same area, we generated all the three types of axes from the subpopulations. Briefly, each population from S1, PPC-RL, and PPC-A was randomly divided into two subpopulations with similar number of neurons. Then, to reduce the computational challenge and to ensure a stable CCA computation, we reduced the dimensionality of each subpopulation with PCA. To ensure that each subpopulation had a similar number of variables, especially for the inter-area axis computation, we kept the top 30 PCs for each area. We then computed the three axis types using each subpopulation. We repeated the random split 10 times, resulting in 20 subpopulations for each area. Only datasets with good imaging quality from both areas by manual curation were included in this analysis.

For the encoding axis, we computed the trial average vector for each task variable (for example, texture 1 versus texture 2 trials), and defined the encoding axis as the difference between these two mean vectors. Shuffled distribution for the encoding axis discrimination index was computed by shuffling the input data. For axis similarity, to account for the differences in the variance captured by each PC, we computed a weighted cosine similarity, defined as 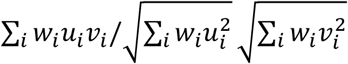 where *u* and *v* are the axes, *i* represents the element in the axis vector, and *w* is the variance explained by the corresponding PC of each element in the axis vector.

### Within-area and inter-area axis analysis

We utilized canonical correlation analysis (CCA) to evaluate the interaction between neuronal populations within and across different areas^23^. CCA works by identifying pairs of dimensions from the neuronal populations in two neuronal populations, optimizing the correlation between their projected activities. For example, given two neuronal populations whose activity is represented by an *n*_x_ × *t* matrix **X** from the first population and a *n*_y_ × *t* matrix **Y** from the second population—where *t* represents the number of time points and *n*_x_ and *n*_y_ are the number of neurons in each respective area— CCA determines min(*n*_x_, *n*_y_) pairs of dimensions, with the correlation between the projections onto these dimensions decreasing from the first pair to the last. Unlike PCA, which independently maximizes the variance explained by the top axes from **X** and **Y**, CCA focuses on maximizing the correlation between the projections of the activity matrices **X** and **Y** onto the identified dimensions.

We performed CCA analysis for the population activity during each task window, where frames pooled across trials were treated as observations. For within-area interaction axes, we performed CCA using the two subpopulations from the same area. For inter-area interaction axes, we performed CCA using two pairs of subpopulations from the two simultaneously imaged areas. We also computed 100 shuffled models by randomizing the trial correspondence between the two subpopulations. We determined the number of significant interaction dimensions using a threshold defined as mean + 1.96 S.D. (standard deviation) of the highest shuffled correlation, from the 100 shuffled models. We also used the shuffled models to generate a shuffled distribution of axis similarity, by computing the weighted cosine similarity with encoding axes. Similarity values from the real models were considered significant if they were higher than the 95% quantile of the shuffled distribution.

### Analysis of correct and incorrect trials

To account for the differences in trial numbers and to ensure enough samples from each trial type over learning (e.g. expert sessions contain more correct trials than incorrect trials), we re-defined sessions in each imaging day to include 50 trials from each trial type (correct and incorrect trials). For each session, we randomly downsampled the trial type with more trials to generate matching number of trials from each trial type. We excluded the sessions with less than 20 trials of either trial type to ensure stable analysis results (final session length: median 50 trials for each trial type, mean±SD 46±6 trials). We repeated this random downsampling procedure 10 times for each dataset, and averaged the results. This procedure was implemented for all analysis concerning comparison between correct and incorrect trials.

### Statistics and Reproducibility

All statistical analysis was done in MATLAB. In general, Wilcoxon signed-rank test was used for paired samples, and Wilcoxon Rank Sum test was used for non-paired samples. No normality test was performed since these tests do not assume normality. Two-sided tests were performed unless otherwise indicated. No statistical methods were used to pre-determine sample sizes, but our sample sizes are similar to those reported in previous publications^1,6,8,26,28,44^. Error bars represent mean±SEM. P-values were corrected for false discovery rate (FDR)<0.05 for multiple testing, using MATLAB function mafdr. Mice were randomly assigned to imaging groups (S1 and PPC-A or S1 and PPC-RL imaging). Stimulus presentation during imaging was fully randomized. Data collection and analysis were not performed blind to the conditions of the experiments.

## Supplemental Information

**Supplemental Figure 1.**
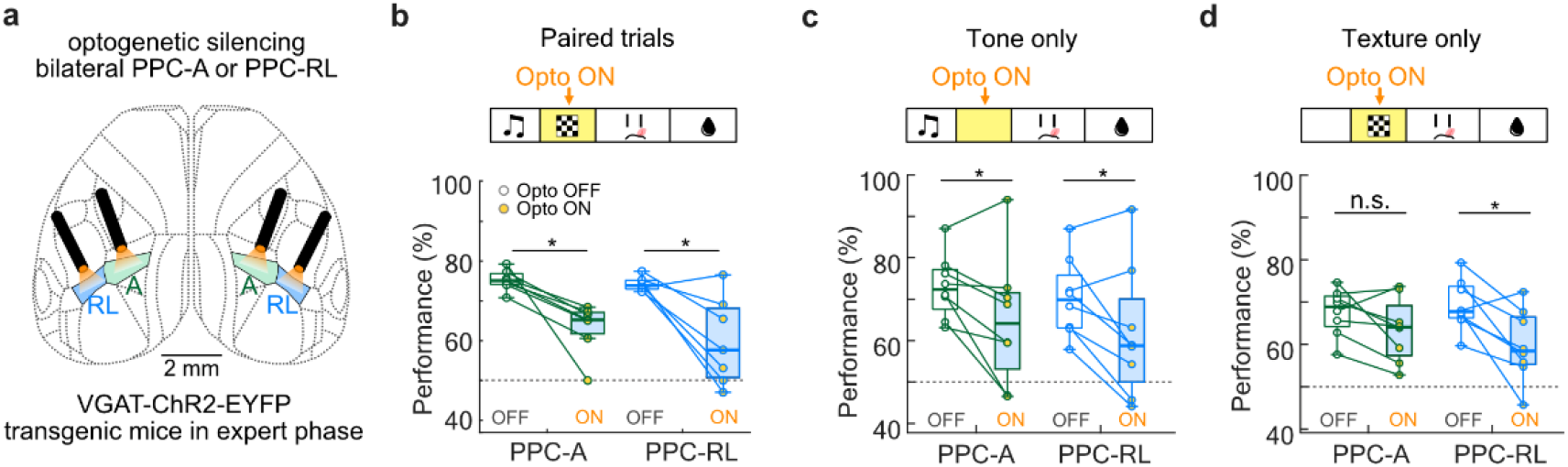
PPC-A and PPC-RL are required for optimal task performance. **(a)** Design of optogenetic inhibition experiment. Optical fibers were bilaterally implanted above PPC-A and PPC-RL of VGAT-ChR2 transgenic mice to allow optogenetic silencing. **(b)** Silencing of either PPC-A or PPC-RL during the texture window reduced task performance. **(c)** Silencing of either PPC-A or PPC-RL during the texture window in tone-only condition reduced task performance. **(d)** Silencing of PPC-RL, but not PPC-A, during the texture window in texture-only condition reduced task performance. (Paired: 7 mice; tone only and texture only: 8 mice).

**Supplemental Figure 2.**
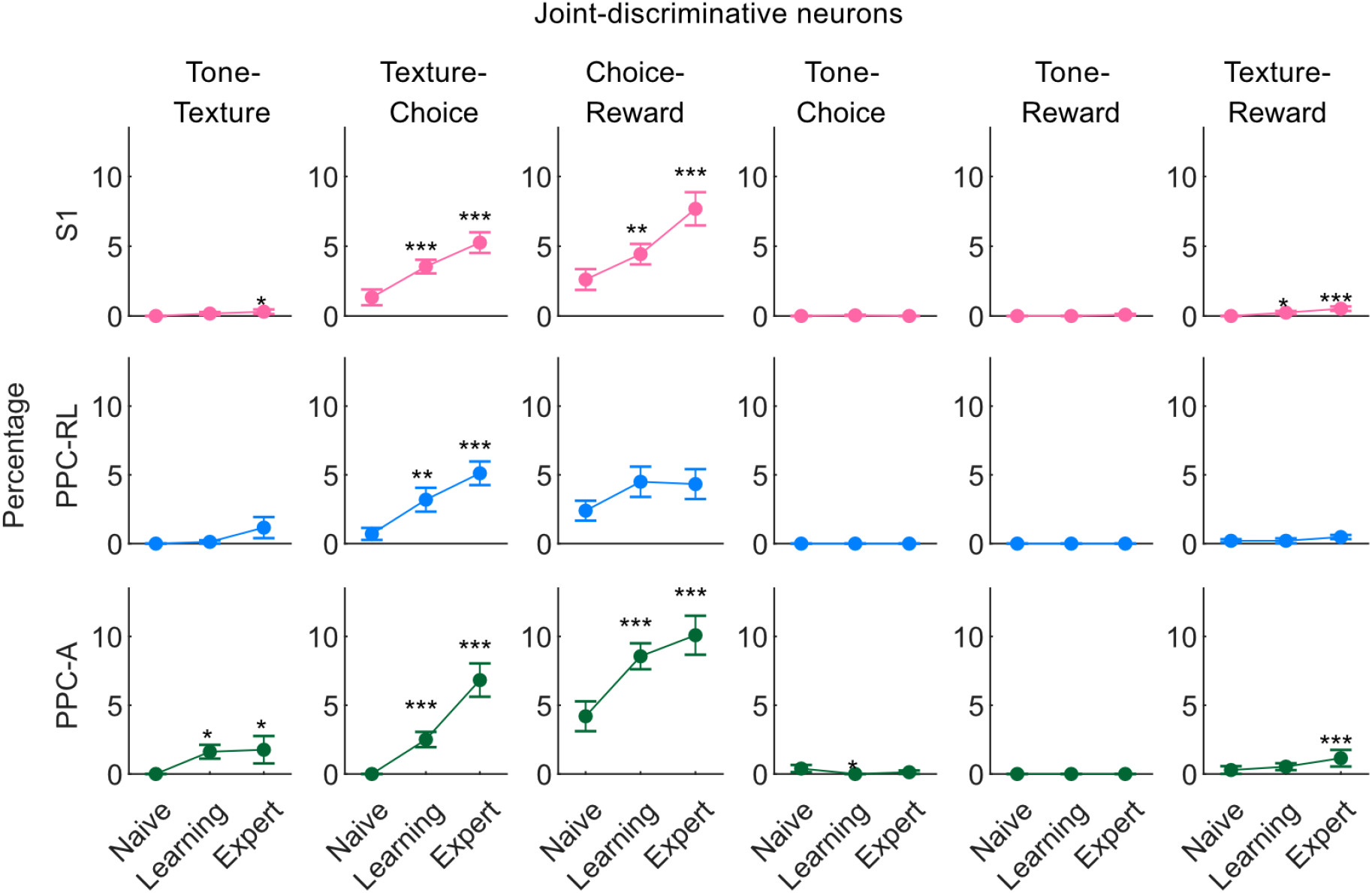
Percentage of joint-discriminative neurons. Percentage of joint-discriminative neurons for pairs of task windows, as indicated on top, for S1 (top row), PPC-RL (middle row), and PPC-A (bottom row). (S1: 13 mice, 130, 188, 149 sessions [naive, learning, expert]; PPC-RL: 13 mice, 70, 77, 104 sessions; PPC-A: 10 mice, 60, 111, 45 sessions; \**p*<0.05, \*\**p*<0.01, \*\*\**p*<0.001; Wilcoxon rank-sum test against naive condition).

**Supplemental Figure 3.**
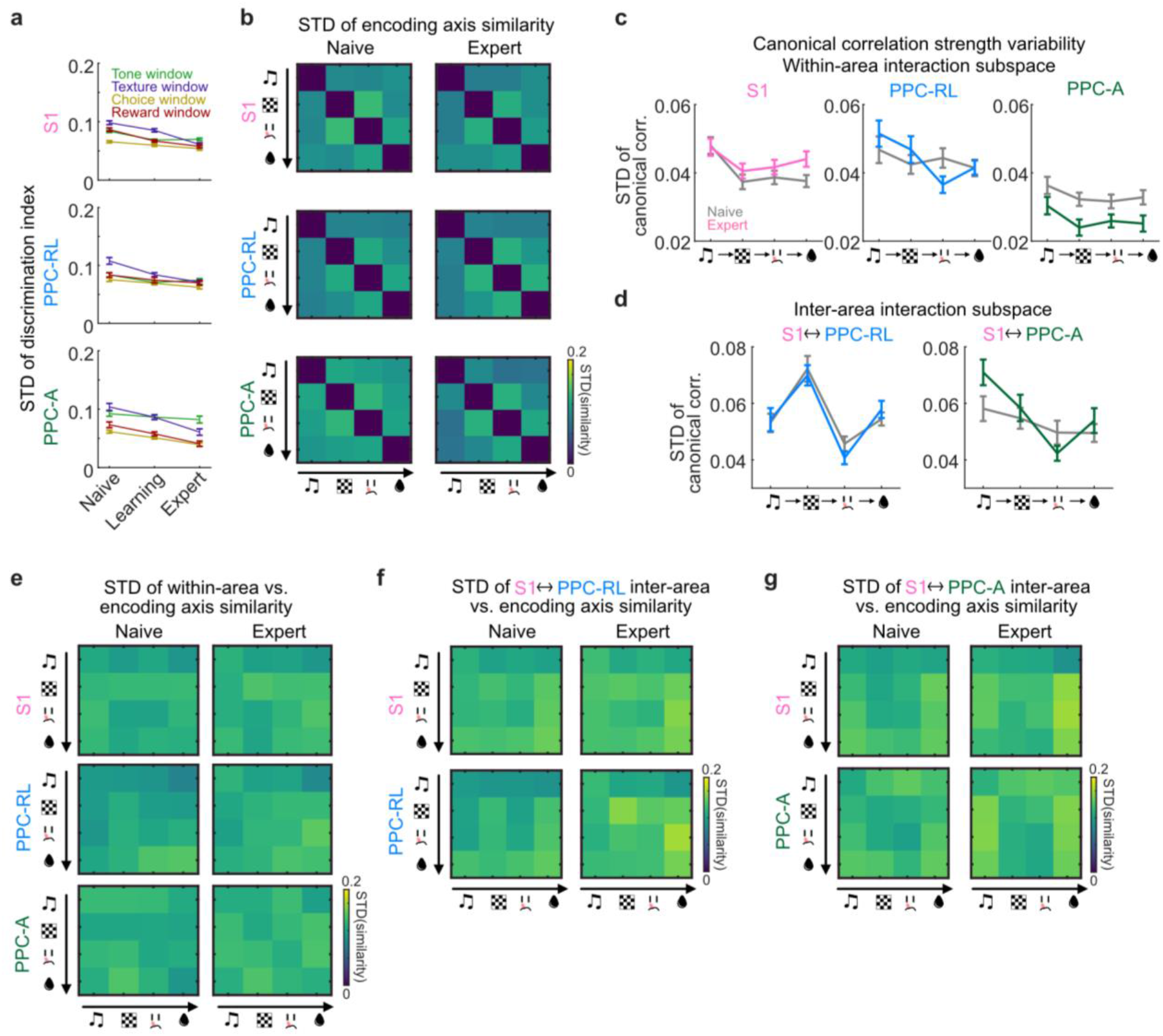
Stability of subpopulation random splits. **(a)** Standard deviation (STD) encoding axis discrimination index in each task window from Fig. 4c, using the 20 subpopulations from 10 random splits in each session. **(b)** STD of encoding axis pairwise similarity of random splits. **(c)** STD of CCA within-area correlation strength of random splits. **(d)** STD of CCA inter-area correlation strength of random splits. **(e)** STD of within-area axis similarity with encoding axis of random splits. **(f)** STD of S1 and PPC-RL inter-area axis similarity with encoding axis of random splits. **(g)** STD of S1 and PPC-A inter-area axis similarity with encoding axis of random splits. (S1: 13 mice, 122, 171, 119 sessions [naive, learning, expert]; PPC-RL: 13 mice, 66, 72, 104 sessions; PPC-A: 10 mice, 56, 99, 33 sessions; S1 and PPC-RL interaction: 13 mice, 66, 72, 86 sessions; S1 and PPC-A interaction: 10 mice, 56, 99, 33 sessions; 10 random splits into 20 subpopulations per session; mean±SEM).

**Supplemental Figure 4.**
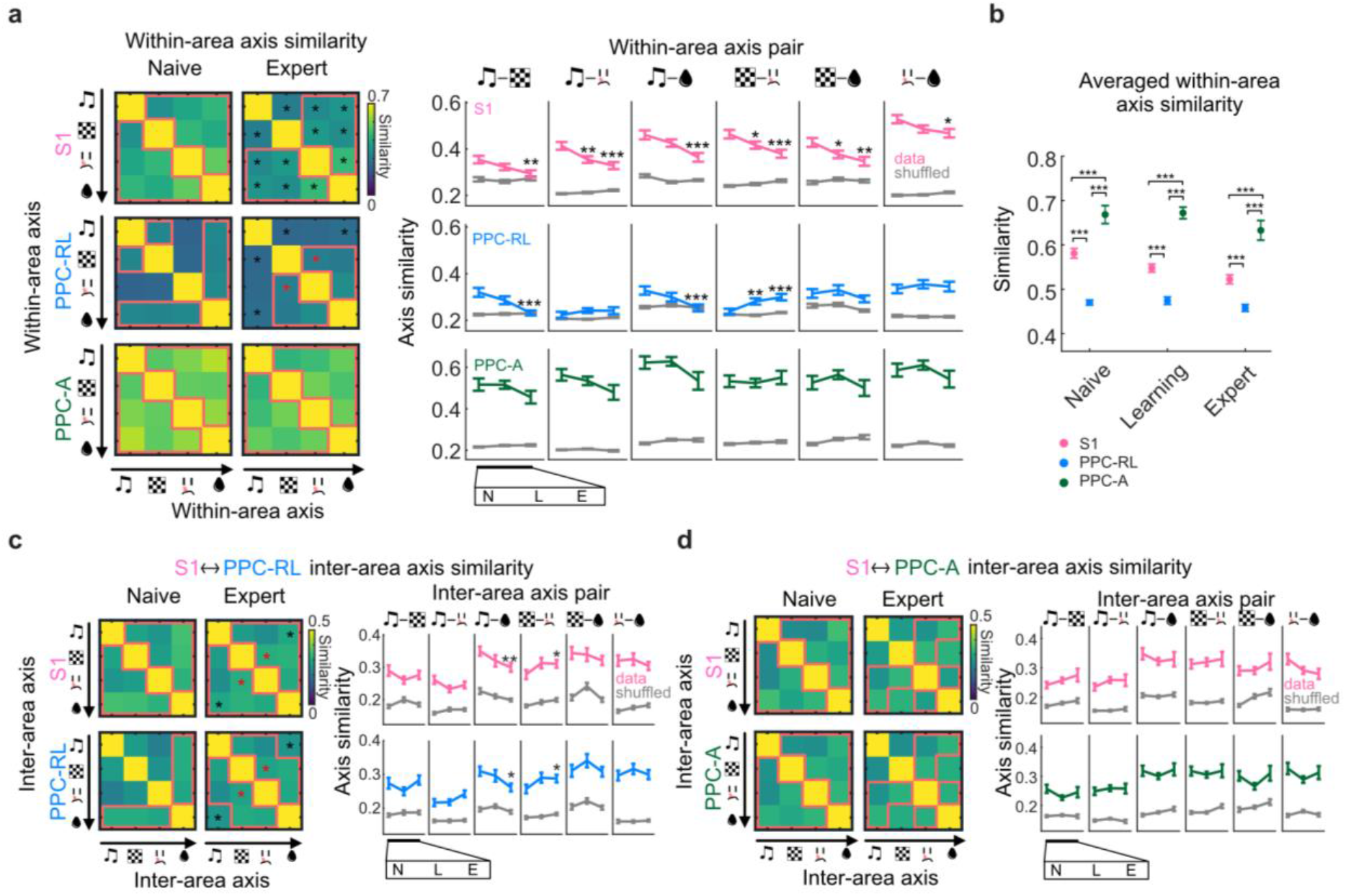
Within-area axis similarity and inter-area axis similarity. **(a)** Within-area axis pairwise similarity. Left: pairwise similarity of within-area axis across task windows. Red lines indicate that the data is above the corresponding shuffled distribution; asterisks indicate that the data from expert condition is significantly higher (red) or lower (black) than the naive condition. Right: quantification of similarity. **(b)** Average within-area axis similarity for each area across learning. **(c)** Inter-area axis pairwise similarity for S1 (top) and PPC-RL (bottom). Left: pairwise similarity of inter-area axis across task windows. Right: quantification of similarity. **(d)** Inter-area axis pairwise similarity for S1 (top) and PPC-A (bottom). Left: pairwise similarity of inter-area axis across task windows. Right: quantification of similarity. (S1: 13 mice, 122, 171, 119 sessions [naive, learning, expert]; PPC-RL: 13 mice, 66, 72, 104 sessions; PPC-A: 10 mice, 56, 99, 33 sessions; S1 and PPC-RL interaction: 13 mice, 66, 72, 86 sessions; S1 and PPC-A interaction: 10 mice, 56, 99, 33 sessions; \**p*<0.05, \*\**p*<0.01, \*\*\**p*<0.001; **a, c, d**, left panels: \**p*<0.05; Wilcoxon rank-sum test against naive condition; mean±SEM).

**Supplemental Figure 5.**
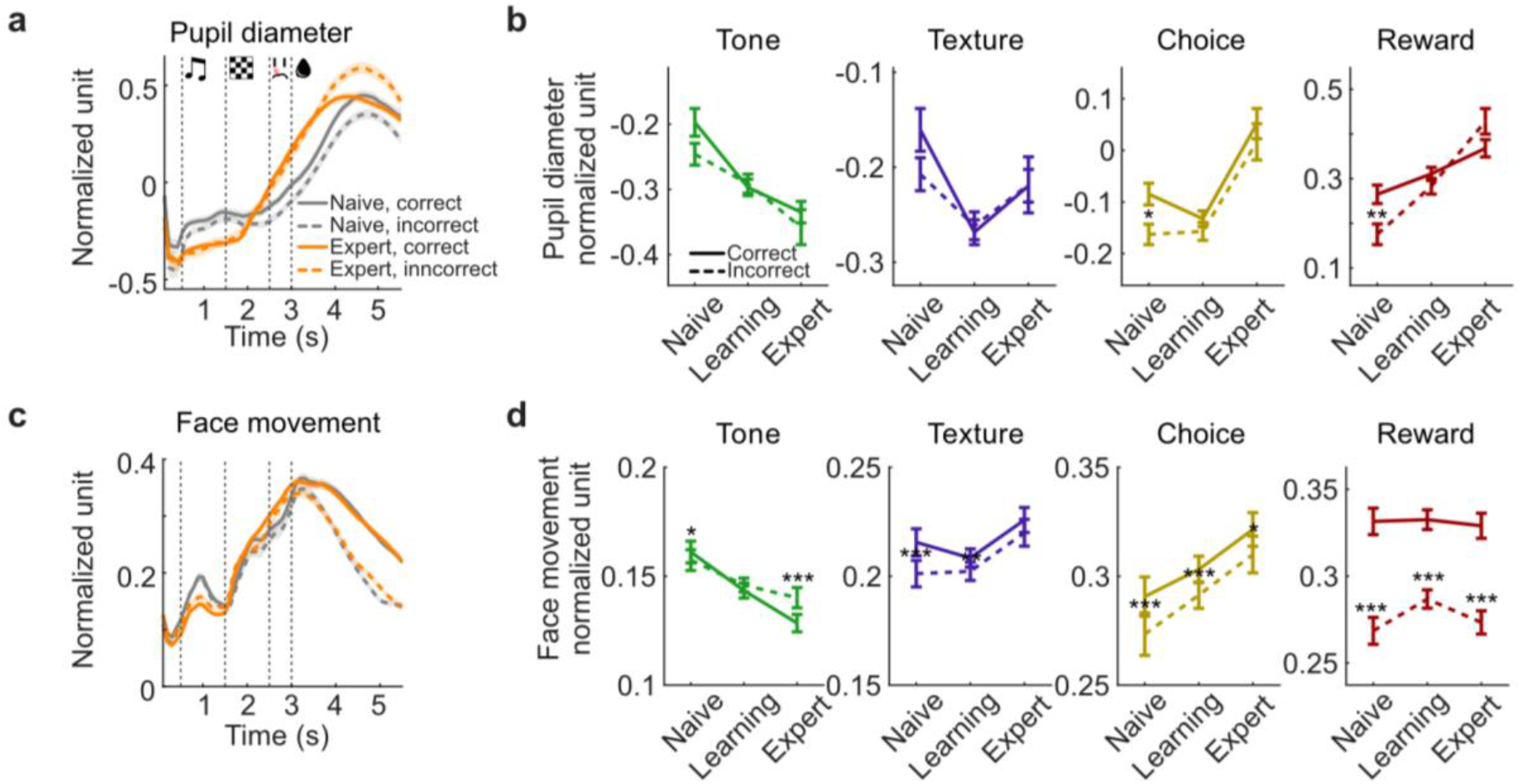
Pupil diameter and face movement during incorrect trials. **(a)** Normalized pupil diameter over trial time, in correct and incorrect trials of naive and expert conditions. **(b)** Quantification pupil diameter in correct and incorrect trials over learning. **(c)** Normalized face movement over trial time, in correct and incorrect trials of naive and expert conditions. **(d)** Quantification face movement in correct and incorrect trials over learning. (16 mice, naive 138 sessions, learning 192 sessions, expert 161 sessions; *p<0.05, **p<0.01, ***p<0.001; Wilcoxon signed rank paired test between correct and incorrect trials; mean±SEM).

**Supplemental Figure 6.**
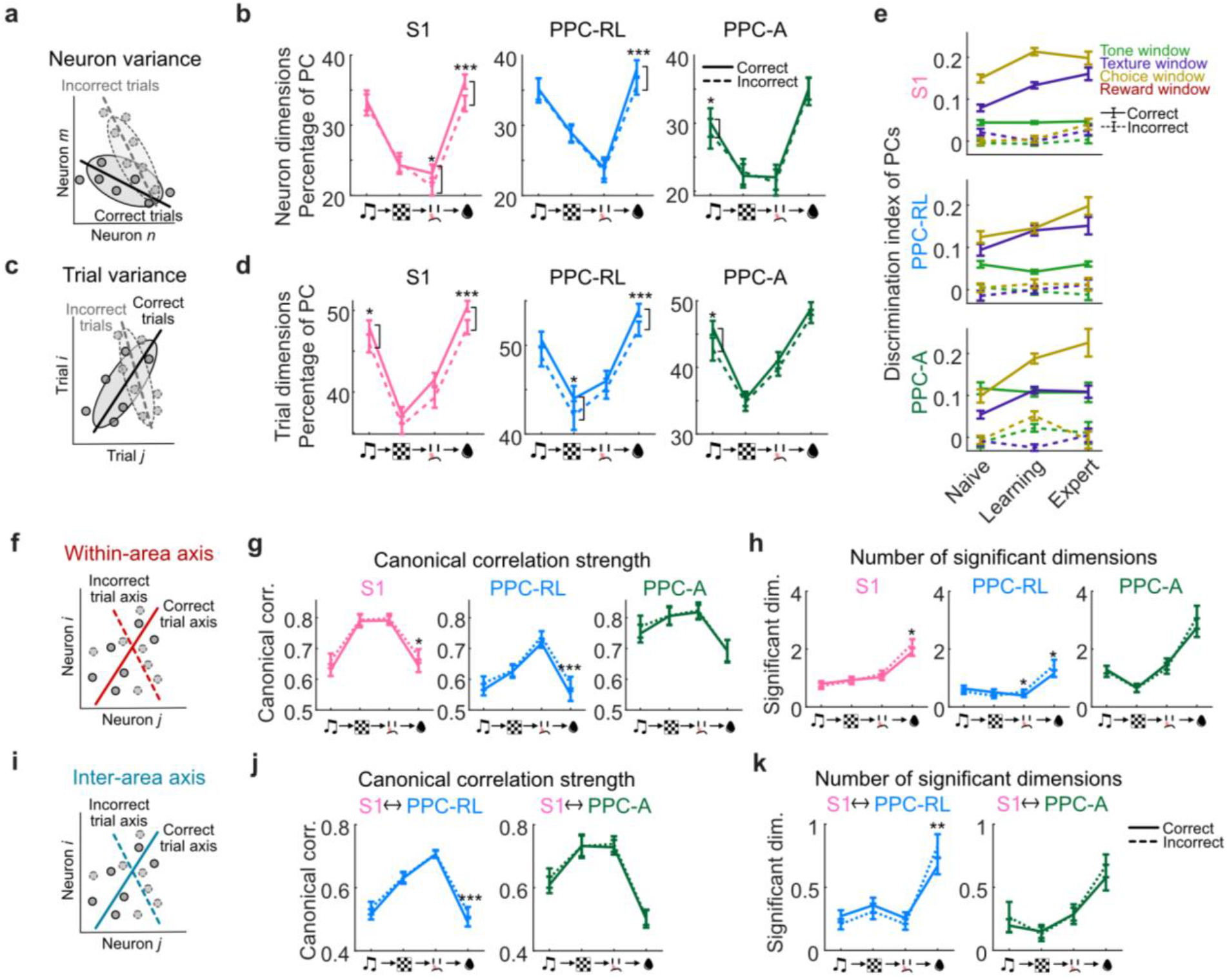
Population dimensionality and interaction subspace. **(a)** Diagram of neuron variance axis in correct and incorrect trials. PCA was performed using only correct or incorrect trials, with neurons as variables. **(b)** Percentage of PCs that explains 70% of variance in neuron space during separate task windows, from correct trials (solid lines) and incorrect trials (dashed lines). **(c)** Diagram of trial variance axis in correct and incorrect trials. PCA was performed using only correct or incorrect trials, with trials as variables. **(d)** Percentage of PCs that explains 70% of variance in trial space, over the trial time, from correct trials (solid lines) and incorrect trials (dashed lines). **(e)** Session-averaged discrimination index of top 5 PCs, during correct trials (solid lines) and incorrect trials (dashed lines). **(f)** Diagram of generating within-area axis from correct and incorrect trials. **(g)** Number of significant dimensions of within-area interactions, in naive and expert condition for each task window in correct and incorrect trials. **(h)** Inter-area canonical correlation strength for each task window in correct and incorrect trials. **(i)** Diagram of generating inter-area axis from correct and incorrect trials. **(j)** Within-area canonical correlation strength for each task window in correct (solid lines) and incorrect (dashed lines) trials. **(k)** Number of significant dimensions of inter-area interactions for each task window in correct and incorrect trials. (S1: 13 mice, 149, 156, 48 sessions [naive, learning, expert]; PPC-RL: 13 mice, 79, 63, 33 sessions; PPC-A: 10 mice, 70, 93, 15 sessions; S1 and PPC-RL interaction: 13 mice, 79, 63, 33 sessions; S1 and PPC-A interaction: 10 mice, 70, 93, 15 sessions; \**p*<0.05, \*\**p*<0.01, \*\*\**p*<0.001; Wilcoxon signed rank paired test between correct and incorrect trials; mean±SEM).

**Supplemental Figure 7.**
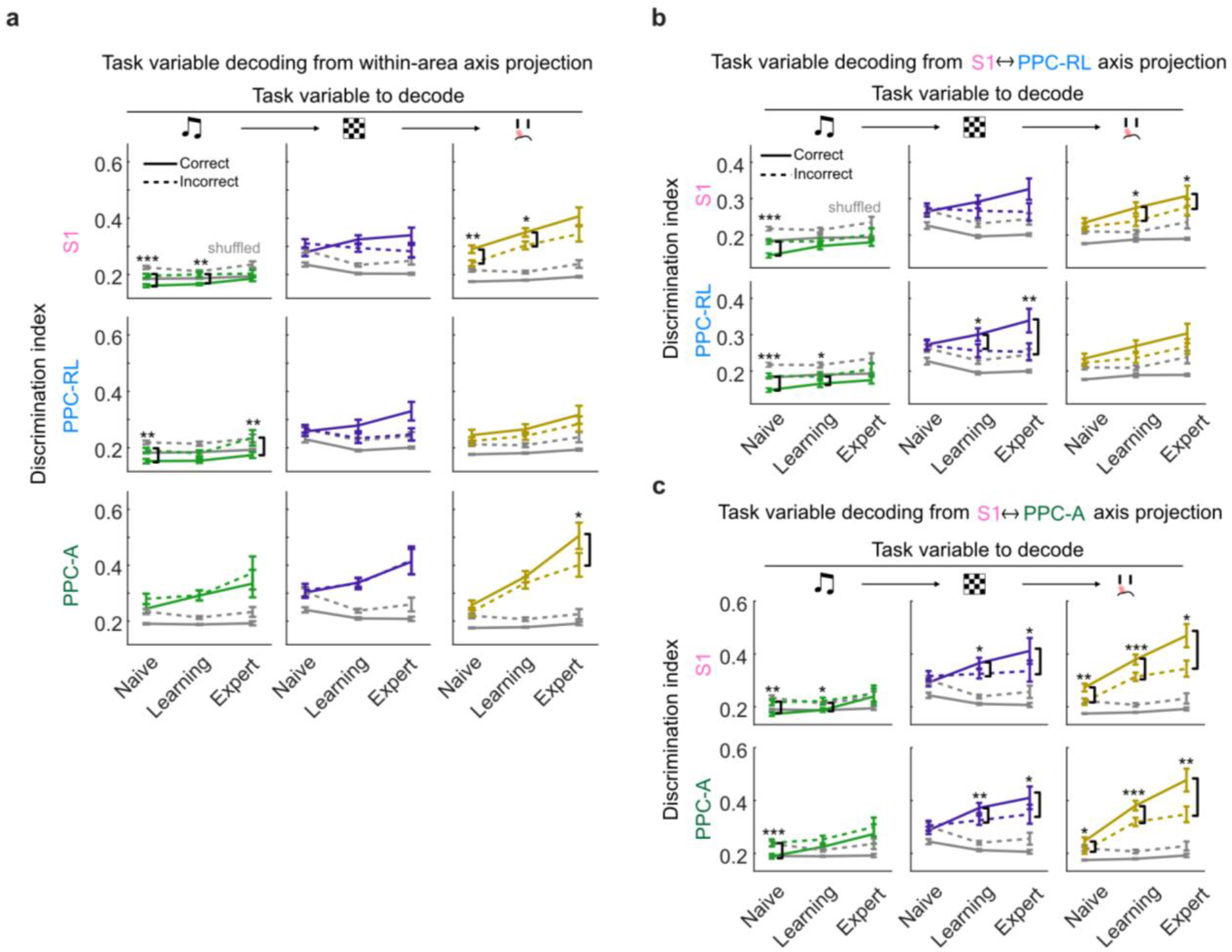
Discrimination index of projected activity on within- and inter-area for correct and incorrect trials. (a) The absolute value of discrimination index of population activity projected onto the within-area axes for S1 (top row), PPC-RL (middle), and PPC-A (bottom). Decoding was performed on projections from the corresponding task windows and axes (e.g., decoding texture using texture window activity projected onto the within-area axis of texture window). Solid lines represent correct trials, dashed lines represent incorrect trials; gray lines represent 95% quantile of shuffled data. (b) The absolute value of discrimination index of population activity projected onto the S1 and PPC-RL inter-area axes. (c) The absolute value of discrimination index of population activity projected onto the S1 and PPC-A inter-area axes. (S1: 13 mice, 149, 156, 48 sessions [naive, learning, expert]; PPC-RL: 13 mice, 79, 63, 33 sessions; PPC-A: 10 mice, 70, 93, 15 sessions; S1 and PPC-RL interaction: 13 mice, 79, 63, 33 sessions; S1 and PPC-A interaction: 10 mice, 70, 93, 15 sessions;*p<0.05, **p<0.01, ***p<0.001; Wilcoxon signed rank paired test between correct and incorrect trials; mean±SEM).

## References

1. Poort, J. et al. Learning Enhances Sensory and Multiple Non-sensory Representations in Primary Visual Cortex. Neuron 86, 1478–1490 (2015).

2. Peron, S. P. P. et al. A Cellular Resolution Map of Barrel Cortex Activity during Tactile Behavior. Neuron 86, 783–799 (2015).

3. Peters, A. J., Chen, S. X. & Komiyama, T. Emergence of reproducible spatiotemporal activity during motor learning. Nature 510, 263–267 (2014).

4. Bale, M. R. et al. Sequence Learning Induces Selectivity to Multiple Task Parameters in Mouse Somatosensory Cortex. Curr. Biol. 31, 1–13 (2020).

5. Huber, D. et al. Multiple dynamic representations in the motor cortex during sensorimotor learning. Nature 484, 473–478 (2012).

6. Chen, J. L. et al. Pathway-specific reorganization of projection neurons in somatosensory cortex during learning. Nat. Neurosci. 18, 1101–1108 (2015).

7. Gdalyahu, A. et al. Associative Fear Learning Enhances Sparse Network Coding in Primary Sensory Cortex. Neuron 75, 121–132 (2012).

8. Driscoll, L. N., Pettit, N. L., Minderer, M., Chettih, S. N. & Harvey, C. D. Dynamic Reorganization of Neuronal Activity Patterns in Parietal Cortex. Cell 170, 986-999.e16 (2017).

9. Golub, M. D. et al. Learning by neural reassociation. 21, 607–616 (2018).

10. Failor, S. W., Carandini, M. & Harris, K. D. Visuomotor association orthogonalizes visual cortical population codes. bioRxiv (2022) doi:10.1101/2021.05.23.445338.

11. Chadwick, A. et al. Learning shapes cortical dynamics to enhance integration of relevant sensory input. Neuron 111, 106-120.e10 (2023).

12. Haimerl, C., Ruff, D. A., Cohen, M. R., Savin, C. & Simoncelli, E. P. Targeted V1 comodulation supports task-adaptive sensory decisions. Nat. Commun. 2023 141 14, 1–15 (2023).

13. Kaufman, M. T., Churchland, M. M., Ryu, S. I. & Shenoy, K. V. Cortical activity in the null space: permitting preparation without movement. Nat. Neurosci. 2014 173 17, 440–448 (2014).

14. Yan, Y. et al. Perceptual training continuously refines neuronal population codes in primary visual cortex. Nat. Neurosci. 2014 1710 17, 1380–1387 (2014).

15. Runyan, C. A., Piasini, E., Panzeri, S. & Harvey, C. D. Distinct timescales of population coding across cortex. Nature 548, 92–96 (2017).

16. Sadtler, P. T. et al. Neural constraints on learning. Nature 512, 423–426 (2014).

17. Hennig, J. A. et al. Learning is shaped by abrupt changes in neural engagement. Nat. Neurosci. 24, 727–736 (2021).

18. Makino, H. & Komiyama, T. Learning enhances the relative impact of top-down processing in the visual cortex. Nat. Neurosci. 18, 1116–1122 (2015).

19. Le Merre, P. et al. Reward-Based Learning Drives Rapid Sensory Signals in Medial Prefrontal Cortex and Dorsal Hippocampus Necessary for Goal-Directed Behavior. Neuron 97, 83-91.e5 (2018).

20. Athalye, V. R., Ganguly, K., Costa, R. M. & Carmena, J. M. Emergence of Coordinated Neural Dynamics Underlies Neuroprosthetic Learning and Skillful Control. Neuron 93, 955-970.e5 (2017).

21. Bondy, A. G., Haefner, R. M. & Cumming, B. G. Feedback determines the structure of correlated variability in primary visual cortex. Nat. Neurosci. 2018 214 21, 598–606 (2018).

22. Yu, Y., Stirman, J. N., Dorsett, C. R. & Smith, S. L. Visual information is broadcast among cortical areas in discrete channels. bioRxiv 469114 (2024) doi:10.1101/469114.

23. Han, S. & Helmchen, F. Behavior-relevant top-down cross-modal predictions in mouse neocortex. Nat. Neurosci. 27, 298–308 (2024).

24. Chen, J. L., Voigt, F. F., Javadzadeh, M., Krueppel, R. & Helmchen, F. Long-range population dynamics of anatomically defined neocortical networks. Elife 5, (2016).

25. Lyamzin, D. & Benucci, A. The mouse posterior parietal cortex: Anatomy and functions. Neurosci. Res. 140, 14–22 (2019).

26. Gallero-Salas, Y. et al. Sensory and Behavioral Components of Neocortical Signal Flow in Discrimination Tasks with Short-Term Memory. Neuron 109, 1–14 (2020).

27. Harvey, C. D., Coen, P. & Tank, D. W. Choice-specific sequences in parietal cortex during a virtual-navigation decision task. Nature 484, 62–68 (2012).

28. Gilad, A. & Helmchen, F. Spatiotemporal refinement of signal flow through association cortex during learning. Nat. Commun. 11, 1–14 (2020).

29. Song, Y. H. et al. A Neural Circuit for Auditory Dominance over Visual Perception. Neuron 93, 940-954.e6 (2017).

30. Gilad, A., Gallero-Salas, Y., Groos, D. & Helmchen, F. Behavioral Strategy Determines Frontal or Posterior Location of Short-Term Memory in Neocortex. Neuron 99, 814-828.e7 (2018).

31. Pachitariu, M. et al. Suite2p: beyond 10,000 neurons with standard two-photon microscopy. bioRxiv (2017) doi:10.1101/061507.

32. Gurnani, H. & Cayco Gajic, N. A. Signatures of task learning in neural representations. Curr. Opin. Neurobiol. 83, 102759 (2023).

33. Pancholi, R., Ryan, L. & Peron, S. Learning in a sensory cortical microstimulation task is associated with elevated representational stability. Nat. Commun. 2023 141 14, 1–14 (2023).

34. Gallego, J. A., Perich, M. G., Miller, L. E. & Solla, S. A. Neural Manifolds for the Control of Movement. Neuron 94, 978–984 (2017).

35. Bernardi, S. et al. The Geometry of Abstraction in the Hippocampus and Prefrontal Cortex. Cell 183, 954-967.e21 (2020).

36. Semedo, J. D., Zandvakili, A., Machens, C. K., Yu, B. M. & Kohn, A. Cortical Areas Interact through a Communication Subspace. Neuron 102, 249-259.e4 (2019).

37. Ebrahimi, S. et al. Emergent reliability in sensory cortical coding and inter-area communication. Nat. 2022 6057911 605, 713–721 (2022).

38. Veuthey, T. L., Derosier, K., Kondapavulur, S. & Ganguly, K. Single-trial cross-area neural population dynamics during long-term skill learning. Nat. Commun. 11, 1–15 (2020).

39. Bartolo, R. & Averbeck, B. B. Prefrontal Cortex Predicts State Switches during Reversal Learning. Neuron 106, 1044-1054.e4 (2020).

40. Tang, E. et al. Effective learning is accompanied by high-dimensional and efficient representations of neural activity. Nat. Neurosci. 2019 226 22, 1000–1009 (2019).

41. Brincat, S. L., Siegel, M., von Nicolai, C. & Miller, E. K. Gradual progression from sensory to task-related processing in cerebral cortex. Proc. Natl. Acad. Sci. U. S. A. 115, E7202–E7211 (2018).

42. Tafazoli, S. et al. Building compositional tasks with shared neural subspaces. bioRxiv (2024) doi:10.1101/2024.01.31.578263.

43. Law, C. T. & Gold, J. I. Neural correlates of perceptual learning in a sensory-motor, but not a sensory, cortical area. Nat. Neurosci. 2008 114 11, 505–513 (2008).

44. Valente, M. et al. Correlations enhance the behavioral readout of neural population activity in association cortex. 24, 975–986 (2021).

45. Srinath, R., Ruff, D. A. & Cohen, M. R. Attention improves information flow between neuronal populations without changing the communication subspace. Curr. Biol. 0, (2021).

46. Javadzadeh, M. & Hofer, S. B. Dynamic causal communication channels between neocortical areas. Neuron 2470-2483.e7 (2022).

47. Oby, E. R. et al. Dynamical constraints on neural population activity. bioRxiv 2024.01.03.573543 (2024) doi:10.1101/2024.01.03.573543.

48. Kwon, S. E., Yang, H., Minamisawa, G. & O’Connor, D. H. Sensory and decision-related activity propagate in a cortical feedback loop during touch perception. Nat. Neurosci. 2016 199 19, 1243–1249 (2016).

49. Arlt, C. et al. Cognitive experience alters cortical involvement in goal-directed navigation. Elife 11, (2022).

50. Zhuang, J. et al. An extended retinotopic map of mouse cortex. Elife 6, (2017).

51. Harris, J. A. et al. Hierarchical organization of cortical and thalamic connectivity. Nature 575, 195–202 (2019).

